# Semantic and Prosodic Threat Processing in Trait Anxiety

**DOI:** 10.1101/2020.01.24.918375

**Authors:** Simon Busch-Moreno, Jyrki Tuomainen, David Vinson

**Author notes:** Corresponding author; address: room 406, 26 Bedford Way, WC1E 6BT. Please note that the current version of this preprint has substantially changed, as we have corrected an important statistical flaw from previous versions. This has importantly altered results, hence the title change. Due to this flaw, the published manuscript of this study has been withdrawn.

## Abstract

The present study attempts to identify how trait anxiety, measured as worry-level, affects the processing of threatening speech. Two experiments using dichotic listening tasks were implemented; where participants had to identify sentences that convey threat through three different information channels: prosody-only, semantic-only and both semantic and prosody (congruent threat). We expected different ear advantages (left or right) depending on task demands, information type, and worry level. We used a full Bayesian approach for statistical modelling and analysis. Results indicate that when participants made delayed responses (Experiment 1), reaction times barely increased as a function of worry level, but under time pressure (Experiment 2) worry level induced clear decreases in reaction times. We explain these results in terms of multistep models of anxiety and language, concluding that present results mainly indicate effects threat aversion-related over-attention to threat, and that we do not provide enough evidence for supporting the integration of anxiety and language multiphasic models.

## Introduction

Humans can convey emotion information through different channels, and in the particular case of language the manipulation of tone and/or meaning (i.e. prosody and semantics) are common ways to do so. While prosodic information relies on suprasegmental variation of intensity, pitch, voice quality and duration, semantic information relies on segmental information: morphemes (minimal meaningful language units) composed of varying combinations of phonological segments (Liu et al., 2013). These different informational features (suprasegmental information associated with prosody, and segmental information associated with semantics) can develop together in a complex language emission such as an emotional sentence, and can convey emotional information simultaneously (Nygaard et al., 2009; Schirmer and Kotz, 2003). To our knowledge, whether intrinsic affect differences between individuals (e.g. variation in trait anxiety) has differentiable effects on prosody and semantics remains an unexplored problem in language perception and comprehension research. Investigating this possible connection can bring to light possible effects of anxiety on language processing, moving forward the understanding of individual differences in language processing but also refining understanding of speech information properties.

The present study aims to understand the effect of trait anxiety on these information properties of speech. We use dichotic listening (DL) which provides a robust test of functional hemispheric lateralization (Hugdahl, 2011), tapping into features of both speech (language) and anxiety (affect) processing. DL can provide a behavioural test of laterality in such a way that information- and affect-related aspects of processing can be disentangled. Normally, responses to DL tasks that do not involve prosody or emotion indicate a right ear advantage (REA): faster response times and/or higher accuracy for language processing at stimuli presented at the right ear, (Hugdahl, 2011). Differently, DL responses to emotional and/or prosodic stimuli show either diminished REAs or a left ear advantage (LEA) (Godfrey and Grimshaw, 2015; Grimshaw et al, 2003).

The idea of exploring laterality in this way is based on previous theoretical models and supporting evidence indicating that brain hemispheres have different processing functions for both speech’s information features and intrinsic affect. On the language side, evidence suggests the left lateralization of segmental aspects of speech and right lateralization of suprasegmental features of speech (Poeppel, 2003; Poeppel et al., 2007; Zatorre, 2001; Zatorre et al., 2002). On the affect side, the relationship between affect and cognition in anxiety is understood to be mediated by right-lateral prefrontal cortex (Gable et al., 2019). Other approaches distinguish between anxious arousal (physiological hyperarousal) and anxious apprehension (worry), where the first is posited as right lateralized and the second as left lateralized or bilateral (Heller et al., 1997; Nitschke et al., 1999; Spielberg et al., 2013). This could imply that intrinsic lateralization patterns induced by individual differences (e.g. anxiety) could match emotional speech’s lateralization patterns. Hence, different information properties (i.e. semantics or prosody) conveying similar emotions (i.e. threat) could affect anxious people in different ways by enhancing or dampening their inherent lateralization patterns when processing emotional stimuli. This motivates the question: what is the difference between semantic and prosodic comprehension in natural emotional expression as processed by anxious people? Before answering this question, we need to find out the points of connection between speech, emotional language and anxiety processing, if any.

### Emotional Language Lateralization

Neuroscientific research has observed that information conveyed through prosody or semantics/syntax is processed via differently lateralized brain routes (Belin et al, 2004). Other findings indicate that this difference might be due mainly to the emotional content of language stimuli (Liebenthal et al., 2005). These differences, however, may not be exclusive. Indeed, if emotional language lateralization is considered as a phasic process, then differences in lateralization might change at any point of the processing time-course (Schirmer and Kotz, 2006). Hence, some of these differences might be related to informational processing and others to the processing of affect/cognition. Therefore, hemisphericity patterns might be due to both emotional and speech processing, but a particular observed left, right or bilateral orientation might be evident depending on the observed time phase. One model addressing this issue is the multistep model of emotional language, which proposes three main processing stages: early stage perceptual processing, mid stage recognition processing, late stage evaluation processing (Kotz and Paulmann, 2011).

Under this model, early stages involve the processing of acoustic properties (purely acoustic information), where greater right hemisphere (RH) engagement would be associated to prosodic processing and left hemisphere engagement (LH) would be associated to phonological processing (Poeppel, 2003; Zatorre, 2001). This leads to the interpretation that LH might process segmental (phonologically composed words) information better, while RH privileges suprasegmental (prosody) information. Mid stages might involve the emotional recognition of stimuli (e.g. integration of previously processed information), implying greater involvement of RH or LH depending on stimulus type and/or conveyed emotion (Schirmer and Kotz, 2006). Late stages would be associated to informational integration and evaluation of emotional stimuli (Kotz and Paulmann, 2011). Another crucial aspect of the model is that it also considers information transferring between hemispheres (Kotz and Paulmann, 2011). Mechanism of callosal relay have been proposed as important aspects for RH to LH (and vice versa) communication of prosodic and syntactic information (Friederici et al, 2007), and also interhemispheric communication of emotional prosody processing (Ross et al, 1997). In addition, callosal relay mechanisms have been proposed as an explanation for different effects of emotional semantics and prosody processing in a dynamic model of DL (Grimshaw et al, 2003). Hence, the observation of bilateral involvement does not necessarily mean that both hemispheres are processing the same information/task, and the observation of unilateral processing does not necessarily mean that the contralateral hemisphere does not play a role.

With all this in mind, there are some relevant issues that this model does not take into account. First, the process does not need to end at an evaluation stage, as natural responses to emotional stimuli are, in general, behaviourally oriented (Vuilleumier, 2005). Thus, a fourth stage associated to goal-orientation might be required to fully grasp emotional language processing. Goal-orientation can be understood as the interruption or pursuing of an organism’s current goals (Bar-Haim et al., 2007), such as the interruption of current behaviour after the perception of a threatening stimulus in order to re-assess situation and environment. Multistage models of intrinsic affect (i.e. anxiety) have proposed that goal-orientation comprises a fourth stage following three initial stages: pre-attentive evaluation of threat, re-orientation of attention, and threat evaluation (Bar-Haim et al., 2007). This is a remarkable match with models of emotional language processing (Kotz and Paulmann, 2011), which match well with the first three of these stages, characterized in the language literature as: identification, recognition, and evaluation. Hence, after evaluation, a deliberation (goal-orientation) stage is a theoretically relevant (if not necessary) theoretical addition. In this sense, tasks that induce overt behaviour should make such a deliberation stage evident, in which participants need to decide about their responses after evaluating the stimuli. Whether this stage has idiosyncratic lateralization patterns as proposed for the previous three stages (Kotz and Paulmann, 2011) is something that has not been consistently explored yet.

### Anxiety and Threat: Affect Lateralization

The strong effects of anxiety over deliberation processes, such as those induced by worry (Corr and McNaughton, 2012; McLaughlin et al., 2007), could imply that people high in trait anxiety process and respond differently to threatening semantics or prosody, including diversified lateralization patterns. The lateralization of affect might not only depend upon processing the emotional content of a particular stimulus, of any type (not only language), that can induce a specific emotion (e.g. threat-inducing fear or anxiety), but also upon individual differences between participants that may also cause different lateralization patterns. This is especially indicated by studies that demonstrate such variation not only when processing emotional stimuli but also during resting state (Nietschke et al., 2000; Engels et al., 2007).

Variation in lateralization patterns related to anxiety have been discussed from a number of different theoretical perspectives. First, anxiety has been proposed to be elicited by a behavioural inhibition system (BIS), which stops approaching behaviour of the organism in order to allow this organism to scan the environment in search of potential threat (McNaughton and Gray, 2000; Corr and McNaughton, 2012). Second, in the approach-withdrawal model, LH would be more engaged in approach-related emotions, while RH would show more involvement on withdrawal-related emotions (Davidson, 1992). This has been also captured by the valence-arousal model (Heller et al, 1997), where two types of anxiety are distinguished: anxious apprehension (worry-related) and anxious arousal (physiological hyperarousal), processed by LH and RH respectively.

In effect, models of anxiety processing propose BIS as a conflict resolution system (Corr and McNaughton, 2012), where anxiety can be interpreted as a plausible intermediate state between approach and withdrawal, or calm and fear. Here, behaviour inhibition and arousal increase in preparation to approach/withdraw responses when possible or needed (McNaughton and Corr, 2014). In other words, behavioural inhibition for environmental scanning might increase arousal levels (McNaughton and Gray, 2000), which can induce fear-related responses if stimuli within the environment appear threatening enough. Thus, the interplay of lateralization patterns associated to worry and arousal might develop differently through the time-course of stimulus evaluation. Indeed, evidence from a functional magnetic resonance imaging (fMRI) study indicates that emotional language induces different lateralization responses, at different processing stages, for different types of anxiety (Spielberg et al, 2013). Where anxious apprehension is associated to a later and continued involvement of LH structures, interpreted as over-engagement with threat (e.g. rumination), and matching evaluation (mid-phase) and orientation/deliberation (late phase) stages (Bar-Haim et al., 2007). Differently, anxious arousal was associated with a faster and of shorter duration RH response interpreted as over-attention to threat, thus matching pre-attentive (early) and re-orientation (early-mid) stages (Bar-Haim et al., 2007).

Electroencephalography (EEG) research using the event-related potential technique (ERP), which offers high temporal resolution, has observed this over-attention and over-engagement response when threatening faces are used as stimuli (Eldar et al, 2010). Over-engagement with threat has also been observed when people with generalized anxiety disorder respond to threatening images (MacNamara and Hajcak, 2010). Furthermore, recent research has observed that socially anxious people present a right lateralized over-attention response to threatening words (Wabnitz, 2015). Hence, trait anxiety might directly affect the processing of emotional language. Previous EEG evidence indicates that anxiety has an effect on the recognition of prosody (Pell et al, 2015). However, not much is known about the interaction between emotion (threat) as conveyed through different information channels (prosody, semantics) and intrinsic affect (anxiety). If phasic lateralization patterns are integrated in a multistage model of anxiety (Bar-Haim et al, 2007) and this is compared to a multistep model of emotional language (Kotz and Paulmann, 2011), then it might be possible to predict very specific behavioural responses for anxious and non-anxious people. More precisely, there is a possible overlap between language processing mechanisms and anxiety processing mechanisms, which could become evident by comparing how people with higher trait anxiety processes different types of speech (i.e. prosody and semantics) as compared to less anxious people.

### Present Experiment

As previously mentioned, emotional and/or prosodic stimuli show either diminished REAs or a left ear advantage (LEA) in some DL studies (Godfrey and Grimshaw, 2015; Grimshaw et al, 2003), indicating a RH processing preference for emotion and/or prosody. However, few dichotic listening (DL) experiments have researched the effects of anxiety on emotional speech processing (Gadea et al, 2011). They either use speech/prosody as an emotion-eliciting stimulus or use DL mainly as an attentional manipulation technique (Bruder et al., 1999; 2005; Leshem, 2018; Peschard et al., 2016; Sander et al., 2005). As a result, they are limited in the extent to which they reveal the relationship between dynamic variations in emotion language processing (prosody/semantics). Instead, studies focusing on the dynamic properties of emotional language, whether using DL or not (e.g. measuring laterality through electrophysiological measures), do not tend to consider individual differences (e.g. Godfrey and Grimshaw, 2015; Grimshaw et al., 2003; Kotz and Paulmann, 2007; Paulmann and Kotz, 2012; Techentin et al., 2009; Wabacq and Jerger, 2004). Therefore, on one side of the picture speech stimuli are typically treated as generic threatening stimuli, so possible differences induced by the informational features of speech that may vary over time are overlooked. On the other side, participants are typically regarded as a homogeneous group, so possible differences induced by anxiety-related processing, that may vary over time and may differ across informational features are overlooked.

Another important thing to consider is that in natural speech, emotional prosody might not be constrained to a single word, as is the case in the experimental manipulations of most of the studies we have cited above. However, semantics is always constrained by sentence’s structure and lexical meaning. In other words, while a lexical item needs to be identified within a sentence in order for emotional semantics to be recognized, prosody might be expressed from the beginning of a sentence. This makes difficult to generalize from word level, or highly controlled sentences, to real world emotional utterances.

To address these issues, we designed two web-based DL experiments, using semi-naturalistic sentences in order to ensure dynamic language processing beyond the single word level. Participants were asked to discriminate between neutral and threatening sentences (the latter expressing threat via semantics, prosody or both), in a direct-threat condition: identifying whether a threatening stimulus was presented to the left or right ear, and in an indirect-threat condition: identifying whether a neutral stimulus was presented the left or right ear. Participant’s anxiety level was measured by using a psychometric scale. By so doing we were able take advantage of past studies researching the attentional effects of threatening language on anxiety and of studies researching the dynamics of speech’s informational properties within a single study.

Both speech processing and anxiety literature seem to converge on theoretical perspectives incorporating multistep models, so we designed two experiments to tap into different points in processing for which individual variation in anxiety may affect speech. In particular, we aimed to differentiate responses made at late evaluative stages (delayed response) vs. responses made at earlier attentive stages (online response) as early over-attention to threat (Bar-Haim et al., 2007) might affect earlier prosody/semantic lateralization patterns (Kotz and Paulmann, 2011), and later over-engagement with threat (Bar-Haim et al., 2007) might affect later emotional language evaluation stages (Kotz and Paulmann, 2011). Thus, Experiment-1 required participants to wait until after sentences’ offset to respond (delayed response), and Experiment-2 required participants to respond during sentence presentation (online response).

For Experiment-1 we hypothesize that anxious over-engagement with threat at mid-late evaluative stages (Bar-Haim et al., 2007) should increase left hemisphere (LH) engagement (Spielberg et al, 2013), disturbing possible LH to right hemisphere (RH) information transferring (Grimshaw et al., 2003; Kotz and Paulmann, 2011). Hence, we predict that a left ear advantage (LEA), usually observed in DL experiments as an effect of prosody/emotional stimuli (Godfrey and Grimshaw, 2015; Grimshaw et al., 2003), should decrease as a function of anxiety, especially for semantic threat. This implies slower and less accurate responses for anxious people at their left ear when responding to semantically threatening but prosodically neutral stimuli (which we named Semantic stimuli). As present sentences are naturalistic, they have varied durations, but are long on average (∼2s). This implies that answering after sentence’s offset emphasizes late stage processing, understood to start at around 400ms (Kotz and Paulmann, 2011), followed by deliberation (∼600ms). This late stage could be sustained for a long period of time, as it is characterized by a cyclic BIS process (McNaughton el at., 2013). Hence, if trait anxiety extends deliberation through excessive worry, then responses locked to sentence’s offset should be slower.

For Experiment-2 we expect that, as responses are forced to be faster (online), prosody should induce the most noticeable effects, as online responses may overlap with early-mid emotional processing stages (Kotz and Paulmann, 2011). Therefore, we hypothesize that higher anxiety should reduce LH involvement (Spielberg et al., 2013) due to over-attention to threat effects, characteristic of earlier-mid processing stages (Bar-Haim et al., 2007). Hence, we predict an enhanced LEA for highly anxious participants, especially for prosodically threatening but semantically neutral stimuli (which we named Prosody stimuli). Thus, faster and more accurate responses for anxious people at their left ear when attending prosodic stimuli. In other words, as participants are required to answer as fast as possible, and prosody is readily identifiable in each sentences, but semantics required the identification of lexical items, processes before quick responses (∼100, ∼200ms) should take precedence for prosody, while semantics might be affected by later processes (∼400ms) as responses could be naturally slower independent of anxiety.

## Methods

### Experiment 1: Delayed Response

#### Participants

Participants were recruited using Prolific (prolific.ac). Only participants reporting being right-handed, having English as first language, without hearing and neurological/psychiatric disorders, and using only a desktop or laptop to answer the experiment were recruited. After exclusion, due to poor accuracy or not finishing the task properly, 44 participants (mean age = 31.7, 27 females) were retained (26 excluded). Participants were remunerated on a £7.5/hour rate. All participants gave their informed consent before participating. It is important to clarify, the web-based nature of the experiment implies that task compliance levels could be low, as there is no direct control over participants meeting requested requirements (e.g. appropriate headphones) or performance (e.g. answering randomly). For this reason, and also to avoid issue related to possible impulsive behaviour or to age-related audition loss, we decided to accept participants well above the adolescence threshold and amply below critical ages for audition loss. Hence, only participants between 24 and 40 years old were accepted to take part.

#### Materials

Four types of sentences were recorded: Prosody (neutral-semantics and threatening-prosody), Semantic (threatening-semantics and neutral-prosody), Congruent (threatening-semantics and threatening-prosody), and Neutral (neutral-semantics and neutral-prosody). We first extracted semantically threatening sentences from movie subtitles by matching the subtitles them with a list of normed threatening words from the extended Affective Norms for English Words (ANEW) (Warriner et al., 2013). For the present study, any word over 5 points in the arousal scale, and below 5 points in the valence and dominance scales was considered threatening (these scales ranged from 1 to 9 points). Every word with less than 5 arousal points and between 4 and 6 (inclusive) valence points was considered neutral. Words’ frequencies were extracted from SUBTLEX-UK (van Heuven et al., 2014), only sentences containing words with Zipf log frequencies over 3 were included. Before recording, ten participants rated the threat level of each visually presented sentence by using a 0-8 Likert scale presented in Gorilla (gorilla.sc). Sentences’ mean ratings were analysed using the Bayesian Estimation Superseeds t-test (BEST) method (Kruschke, 2013). Threatening semantics’ ratings (m = 5.48) were considerably higher than neutral semantics’ ratings (m = 0.3). See Annex for detailed results.

After this, sentences were recorded in an acoustically isolated chamber using a RODE NT1-A1 microphone by a male English speaker. The speaker was not a professional actor or voice actor (i.e. untrained or naïve speaker). The speaker was instructed to speak in what he considered his own angry threatening/angry or neutral voice for recording Prosody/Congruent and Semantic/Neutral sentences respectively. Sentences were not repeated across type (i.e. each type has a unique set of sentences). Neutral dichotic pairs were also unique across conditions (480 different sentences). Due to a technical problem several sentences were recorded with very low amplitude. Therefore, sentences were normalized and cleaned from noise in Audacity (Audacity Team, 2019, audacity.org). Figure 1 shows oscillograms and spectrograms of four example sentences, Table 1 in the Results section also includes a summary of stimulus properties by condition, and the full set of materials can be downloaded from our Open Science Framework (OSF) repository (link in the Data Statement section).

**Figure 1.**
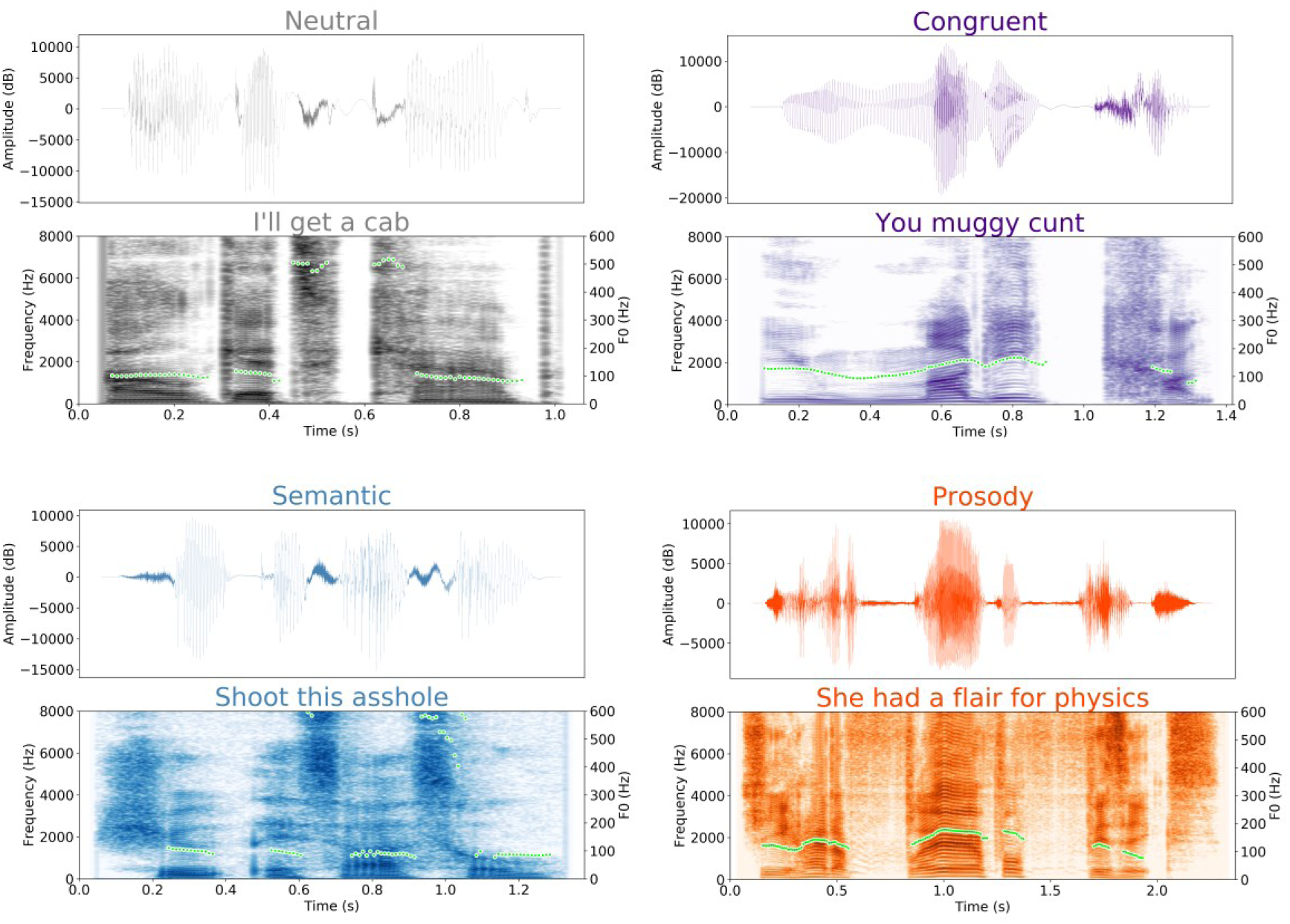
Example of four sentences used as stimuli. Top of each image: oscillogram showing amplitude changes. Bottom of each image: spectrogram showing frequency changes. Top left: neutral prosody and neutral semantics (Neutral). Top right: threatening prosody and threatening semantics (Congruent). Bottom left: neutral prosody and threatening semantics (Semantic). Bottom right: threatening prosody and neutral semantics (Prosody). Green dots indicate fundamental frequency (F0) contours.

**Table 1.**
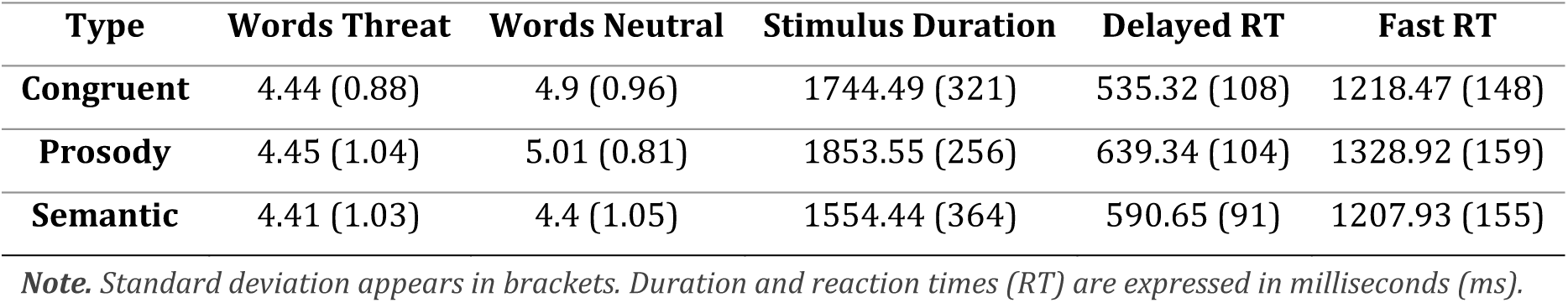
Average number of words, duration and reaction time per stimulus type

Sentences’ average length is 1720.65ms, and acoustic measures were extracted using Parselmouth (Jadoul et al., 2018) Python-Praat interface. Two relevant measures were compared via Bayesian estimation supersedes the t-test (BEST): Median pitch (MP: median F0 across whole sentence) and Hammarberg index (HI: maximum energy differences between the 0-2000hz and 2000-5000hz ranges). While MP is similarly high for Prosody (*m* ≈ 133*Hz*, *SD* ≈ 12*Hz*) and Congruent (*m* ≈ 130*Hz*, *SD* ≈ 9*Hz*) and similarly low for Semantic (*m* ≈ 90*Hz*, *SD* ≈ 2*Hz*) and Neutral (*m* ≈ 96*Hz*, *SD* ≈ 4*Hz*), HI is similarly low for Prosody (*m* ≈ 23*Hz*, *SD* ≈ 4*Hz*) and Congruent (*m* ≈ 24*Hz*, *SD* ≈ 5*Hz*) and similarly high for Semantic (*m* ≈ 32*Hz*, *SD* ≈ 5*Hz*) and Neutral (*m* ≈ 34*Hz*, *SD* ≈ 4*Hz*). Interestingly, semantic ANEW measures show a similar pattern for arousal and valence respectively. Arousal is similarly high for Semantic (*m* ≈ 6.0, *SD* ≈ 0.9) and Congruent (*m* ≈ 5.9, *SD* ≈ 0.9) and similarly low for Prosody (*m* ≈ 3.6, *SD* ≈ 0.9) and Neutral (*m* ≈ 3.9, *SD* ≈ 0.7), and Valence is similarly low for Semantic (*m* ≈ 2.9, *SD* ≈ 1.0) and Congruent (*m* ≈ 3.3, *SD* ≈ 1.2) and similarly high for Prosody (*m* ≈ 5.9, *SD* ≈ 0.8) and Neutral (*m* ≈ 5.8, *SD* ≈ 0.9). See annex for full results of BEST analyses, including ANEW measurements.

This may indicate that as arousal increases and valence decreases, sentences become more semantically threatening, namely convey offense or pain/harm (Borelli et al., 2018; Ho et al., 2015). Similarly, as MP increase (measuring pitch of sentence) and voice HI decreases (measuring voice quality of sentence) the sentence should become more prosodically threatening, express hot anger (Banse and Scherer, 1996; Hammerschmidt and Jürgens, 2007), thus conveying threat. To check this, a random subset of 7 prosody-only sentences was compared to a random subset of 7 neutral sentences in an online rating questionnaire in the same manner as semantic threat. Ten participants rated these spoken sentences in Gorilla (gorilla.sc). Results showed that threatening prosody (m = 5.6, SD = 2.3) is rated as more threatening than neutral prosody (m = 0.71, SD = 1.3). Ratings for semantic threat on written sentences also show that participants (n=22) rate threatening semantics (m = 5.6, SD = 2.2) as more threatening than neutral (m = 0.5, SD = 1.2). An ordered-logistic regression was used to statistically asses these ratings, for results see the Annex.

Next, sentences were paired using Audacity: sentences were paired such as their durations were as similar as possible. Silences between words were extended, never surpassing 40ms, to match sentences’ latencies as closely as possible. After this, sentences were allocated to one of the stereo channels (left or right) of the recording; each pair was copied with mirrored channels. A silence (∼50ms) was placed at the beginning and at end of each pair. This resulted in a total of 480 pairs where 80 sentences of each type (congruent, semantic, prosody) were each paired with a neutral sentence of the same length twice, so every sentence was presented once at each ear.

#### Procedure

Before starting the experiments, participants answered the Penn State Worry Questionnaire (PSWQ) (Meyer et al., 1990) to assess their worry-level, and the Anxious Arousal sub-scale of the Mood and Anxiety Symptoms Questionnaire (MASQ-AA) (Watson et al., 1995) to assess their arousal level. This follows previous approaches (Nitschke et al., 1999), with the difference that we used PSWQ scores as continuous predictor instead of splitting participants between high and low anxiety groups. PSWQ results indicated a distribution which is varied enough in terms of worry level (mean = 47.31, median = 48.0, range [33, 67]). PSWQ measures worry in a scale ranging from 16 to 80 points (median = 48 points), showing a consistent normal distribution in tested samples (mean close to median, as in our samples), and has been shown to have high internal consistence and validity (for details see: Meyer et al., 1990). MASQ-AA scores indicate that participants showed low levels of arousal, as none of them marked above the median. According to previous literature (e.g. Heller et al., 1997; Nitschke et al., 1999; Spielberg et al., 2013), high scores of MASQ-AA would be indicative of trait anxious arousal (hyperarousal), while high scores of PSQW would indicate trait anxious apprehension. Therefore, we can safely assume that our sample does not include participants with high or trait hyperarousal. So, we only included PSWQ scores in the analyses.

After a practice session, participants were randomly assigned to a list containing half of the total number of dichotically paired sentences (threat-neutral pairs) per threatening type (Prosody|Neutral, Semantic|Neutral, Congruent|Neutral), that is 40 pairs per type (120 in total). Sentences’ lists were created previous to the experiment using randomly selected sentences from the total pool. Sentences were presented randomly to participants. In one half of the study they were instructed to indicate at which ear they heard the threatening sentence by pressing the right or left arrow keys (direct-threat condition). In the other half of the study they were instructed to respond in the same way, but indicating which ear they heard the neutral sentence in the dichotic pair (indirect-threat condition). This was intended to address attention effects (Aue et al., 2011; Peschard et al., 2016). Starting ear (left or right) and starting condition (direct- or indirect-threat) were counterbalanced. Participants were told to answer, as fast as possible, only when the sentence finished playing and a bulls-eye (target) image appeared on the screen. A 1400ms inter-stimulus-interval (ISI) was used, and the target image stayed on the screen during this period.

#### Analysis

Reaction time (RT) data were recorded in milliseconds, locked to sentence’s offset. Accuracy was coded as correct=1 and else=0 (including misses and false alarms). Participants with hit rates below 70% were excluded, as lower thresholds are too close to chance. This is mainly due to the nature of web-based experiments, where compliance levels cannot be more directly controlled. Thus, we believe that using too low exclusion criteria (e.g. ∼50% or chance) is not methodologically warranted, as we cannot attest for how much variance or bias is added by unidentified non-compliance. Moreover, by setting a higher criterion for inclusion, we ensure that participants are understanding the sentence content sufficiently for the various proposed stages of processing to occur. Two Bayesian hierarchical models were built for reaction time (RT) and accuracy. The RT model, shown on Figure 2, was the basic model structure for all analyses, based on Kruschke (2015) and Martin’s (2018) guidelines. The model for indirect-threat RT was identical excepting the number of observations (obs. = 5,427). Models for accuracy (Figure 2’s lower panel) also show a different number of observations, where indirect-threat = 5,767 obs. Differences in the number of observations are due to the fact that RT data used only correct responses; also, overlapping responses (those going beyond the ISI), were treated as false alarms.

**Figure 2.**
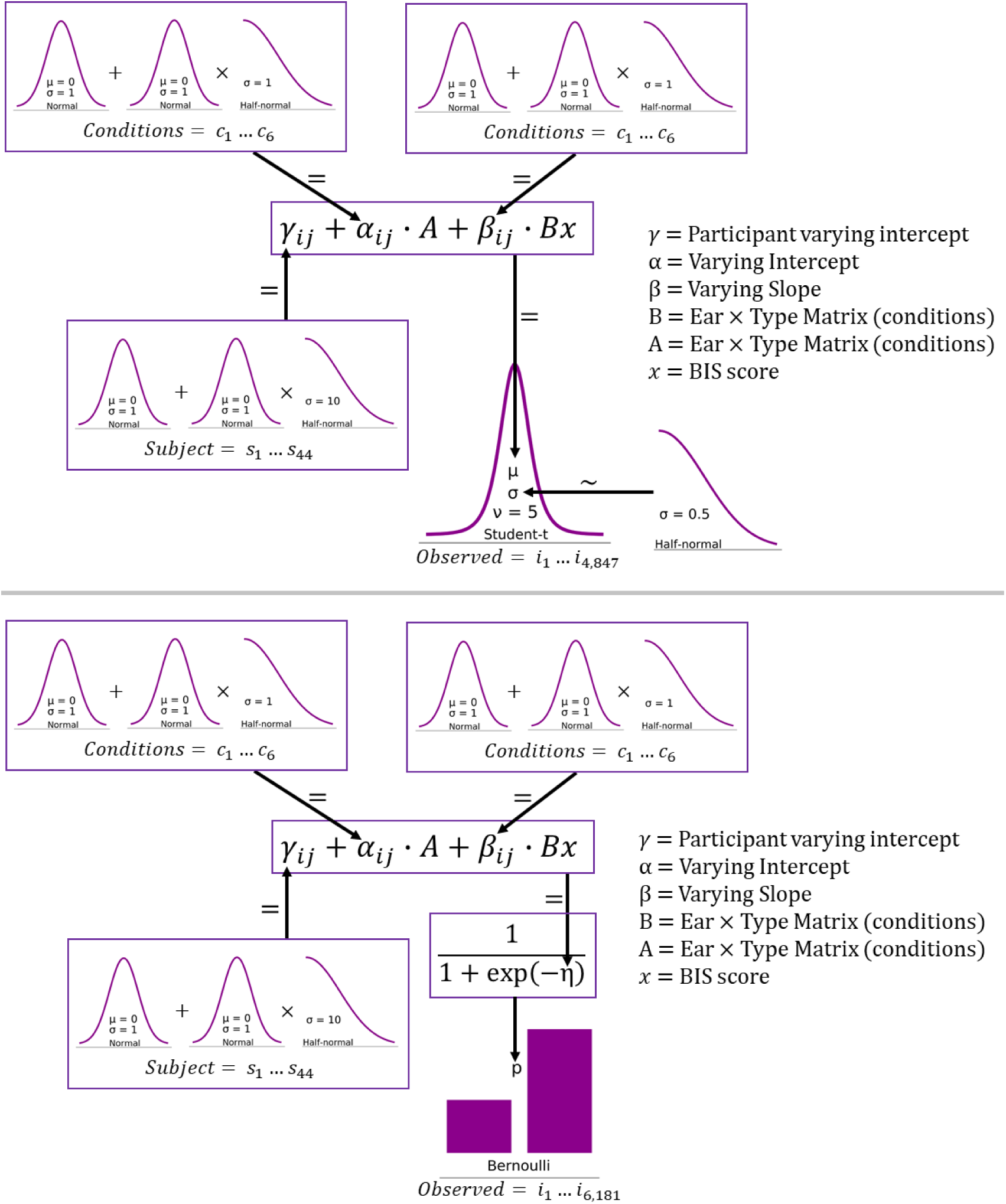
Bayesian models for data analysis. Upper panel: robust regression model for reaction times. Lower panel: logistic regression model for accuracy. Arrows indicate the relationship between a parameter and priors or hyperpriors, where tilde (∼) indicates a stochastic relationship and equal (=) indicates a deterministic relationship. Observations are reaction times (only for correct responses) in milliseconds for the robust regression model, and Bernoulli trials (correct answer =1, else=0) for the logistic regression model. Note that the number of observations and participants correspond to Experiment 1’s (delayed response) direct-threat task. Besides changes in number of observations and participants, models were equivalent for all experiments and tasks.

RT models used a robust regression (Kruschke, 2015) in order to account for outliers through a long-tailed Student-t distribution. In this way, RTs that are implausibly fast or implausibly slow do not need to be removed, but can be dealt with statistically. Both accuracy and RT models were sampled using Markov Chain Monte Carlo (MCMC) No U-turn Sampling (NUTS) as provided by PyMC3 (Salvatier et al., 2016). Four chains of 3000 tuning steps and 2000 samples were used for RT models and four chains of 4000 tuning steps and 4000 samples were used for accuracy models. Plots and model comparisons were produced using Arviz (Kumar et al, 2019) and Matplotlib (Hunter, 2007).

Presently, we are interested in a basic science interpretation of our results rather in an applied science interpretation where threshold decisions are necessary (see Kruschke, 2018). For this reason, we sill focus on the magnitude of effects of regressions’ estimates and on the certainty of these estimates. To account for this, we provide the highest density intervals (HDIs), sometimes referred as credible intervals, for all relevant measures. We interpret overlapping HDIs as less certain effects or no-effects if overlap is wide or total. More information about estimates can found as summary files in our OSF repository (link in the Data Statement section).

### Experiment 2: Fast Response

Experiment 2’s methods were the same as Experiment 1’s methods, same inclusion criteria, same platform, and same materials. Again, the arousal scale did not show any scores above the scale’s median. PSWQ scores indicated a sufficiently varied distribution (mean = 45.22, median = 45.0, range = [26,61]). The only elements that changed from Experiment-1 were the following: 1) As this experiment is understood as more difficult as participants have to answer as fast as possible (before sentence ending) to a widely varied set of sentences, and they are compelled to refrain from any answer as soon as sentences end, accuracy rejection threshold was relaxed to 60% (slightly closer to chance). Given this, 24 participants were excluded and 52 participants (mean age = 31, 24 females) were kept for the final analysis. 2) Participants were instructed to answer, as fast and as accurately as possible, before the sentence finished playing, and to withhold any response when a stop sign image appeared on the screen after sentences’ end. 3) The same robust regression model was applied without duration, which would be inappropriate as participants answer before offset (incomplete sentence’s duration).

## Results

### Experiment 1: Delayed Response

All models sampled properly (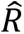 ≤ 1, ESS > 400); energy plots, traceplots and autocorrelation plots also indicate good convergence. Plots and results from these checks, including raw data and full summaries of parameters and conditions, can be found in our OSF repository (link in the Data Statement section).

Reaction time (RT) results from the direct-threat task indicate that worry, as measured by the Penn State Worry Questionnaire (PSWQ), has little to no effect on Congruent sentences or ear, with all regression estimates remaining around 400ms. Prosody sentences at the left ear show a similar pattern, but Prosody at right ear tends to increase around 115ms from the lowest worry score (33 points) to the highest worry score (67 points). Semantic estimates show to be higher overall, indicating slightly slower responses respect to other conditions (around 50 to 80ms), but they show little increase as a function of worry. Note that due to HDI overlap, within and across conditions, these increases cannot be considered certain, where right ear Prosody is the less uncertain increase. Table 2 and Figure 3 summarise these results.

**Figure 3.**
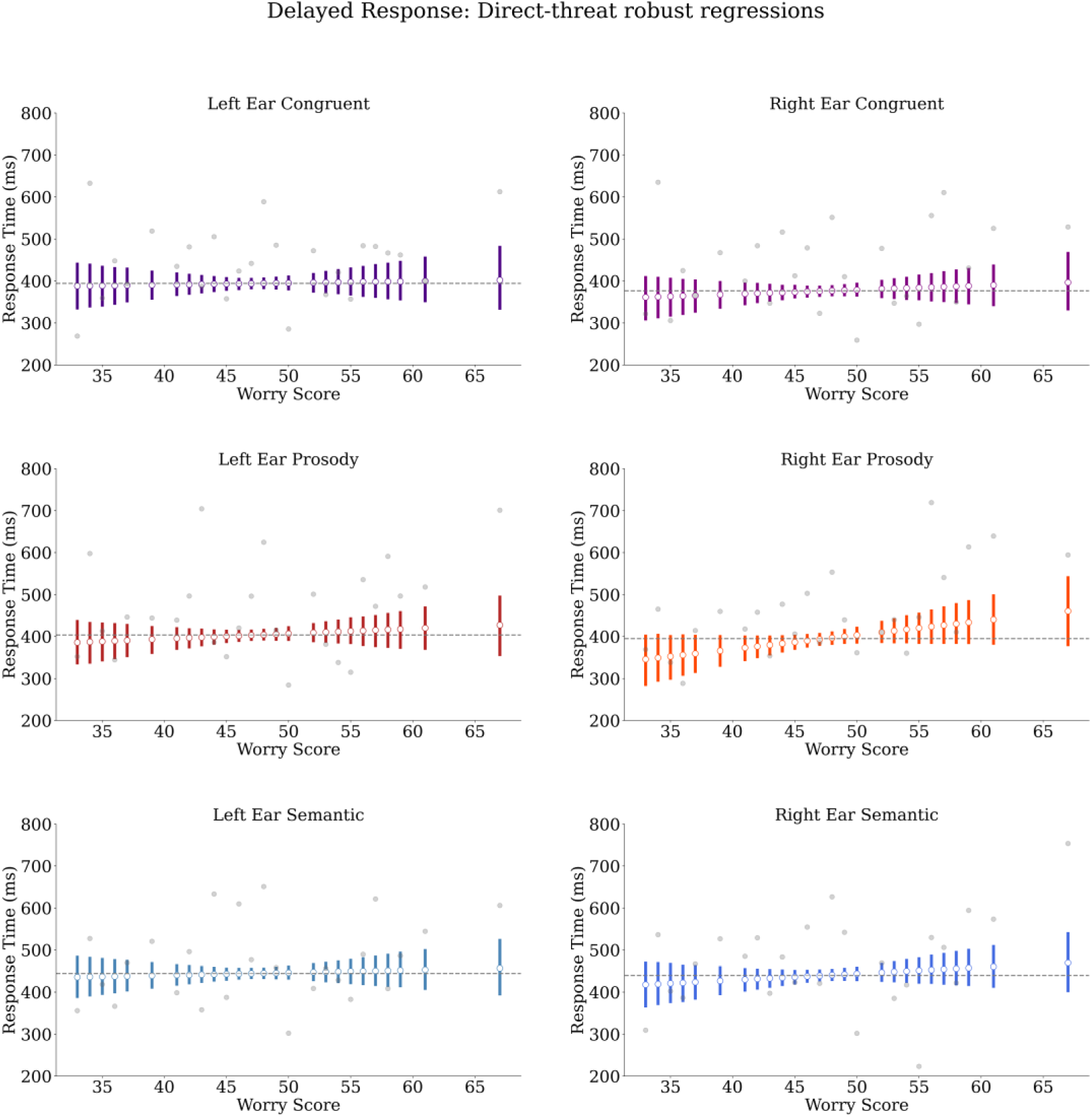
Experiment 1 (delayed response), direct-threat robust regression estimates. White circles indicate posterior means. Coloured bars indicate highest density intervals (HDIs). Grey line indicates estimated worry score median’s posterior median. Grey dots indicate raw means. Note that estimates barely increase as a function of worry, and all conditions indicate HDI overlap, with a reduced overlap only at right ear Prosody.

**Table 2.**
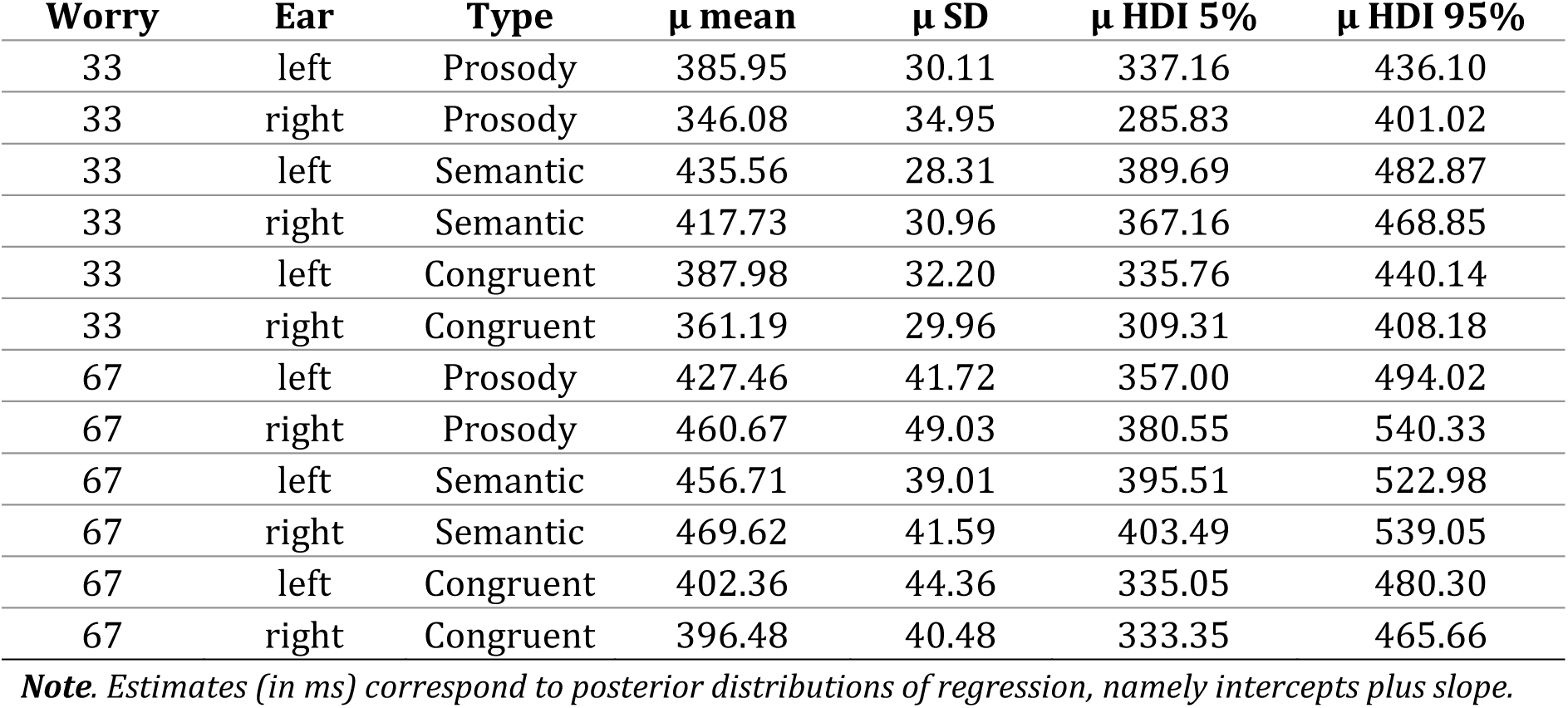
Experiment 1 (delayed response) direct-threat reaction time estimates

In the indirect-threat task, estimates show to be similar, but RT increases seem to be slightly bigger for Congruent. Prosody and semantic conditions also show similar estimates. However, in this case the most certain increase is for Prosody at the left ear that shows the biggest and most certain increase, around 140ms from lowest to highest worry score. See Table 3 and Figure 4 for summaries of these results.

**Figure 4.**
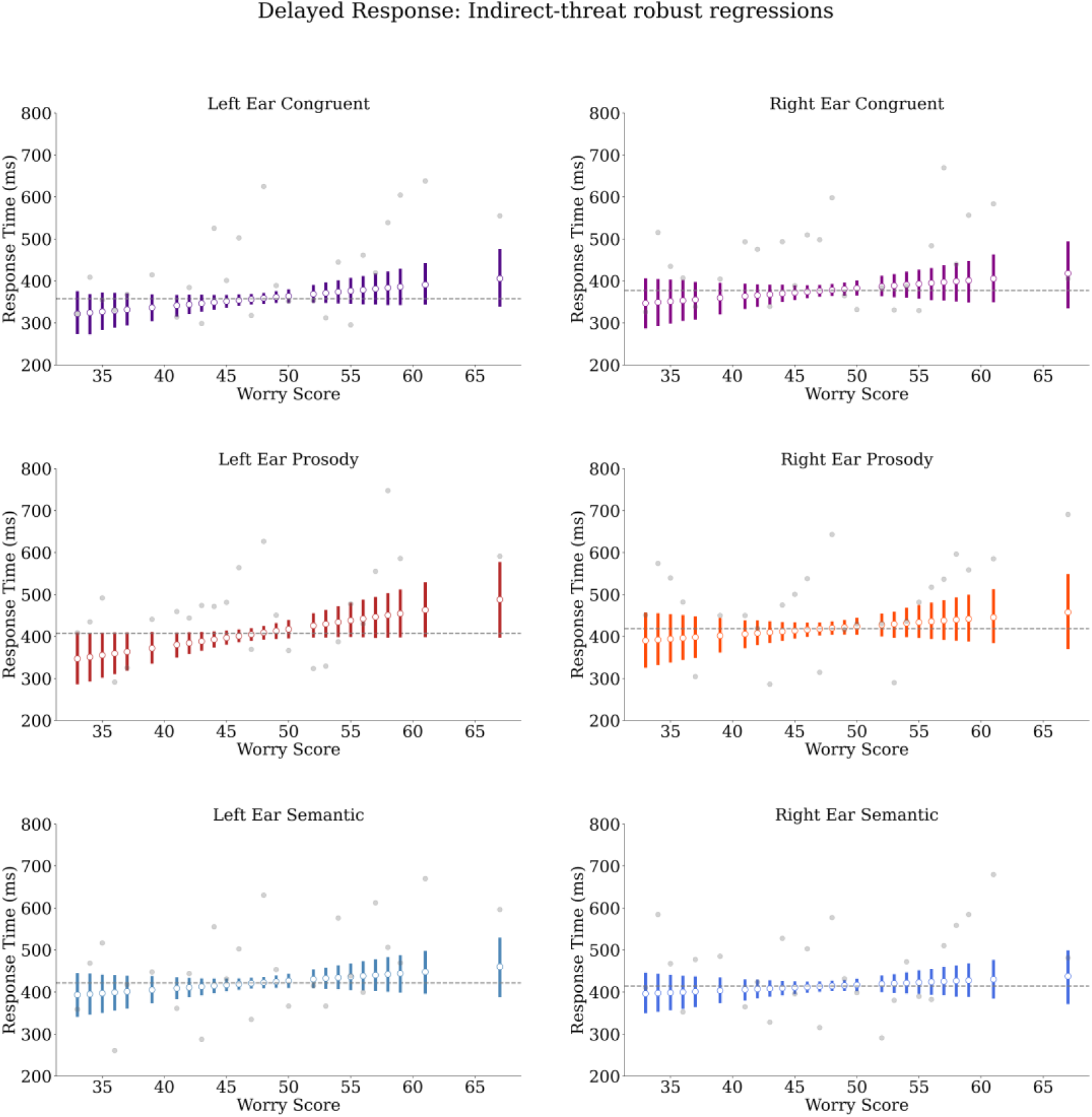
Experiment 1 (delayed response), indirect-threat robust regression estimates. White circles indicate posterior means. Coloured bars indicate highest density intervals (HDIs). Grey line indicates estimated worry score median’s posterior median. Grey dots indicate raw means. Note that estimates barely increase as a function of worry, and all conditions indicate HDI overlap, with very small overlap only at left ear Prosody.

**Table 3.**
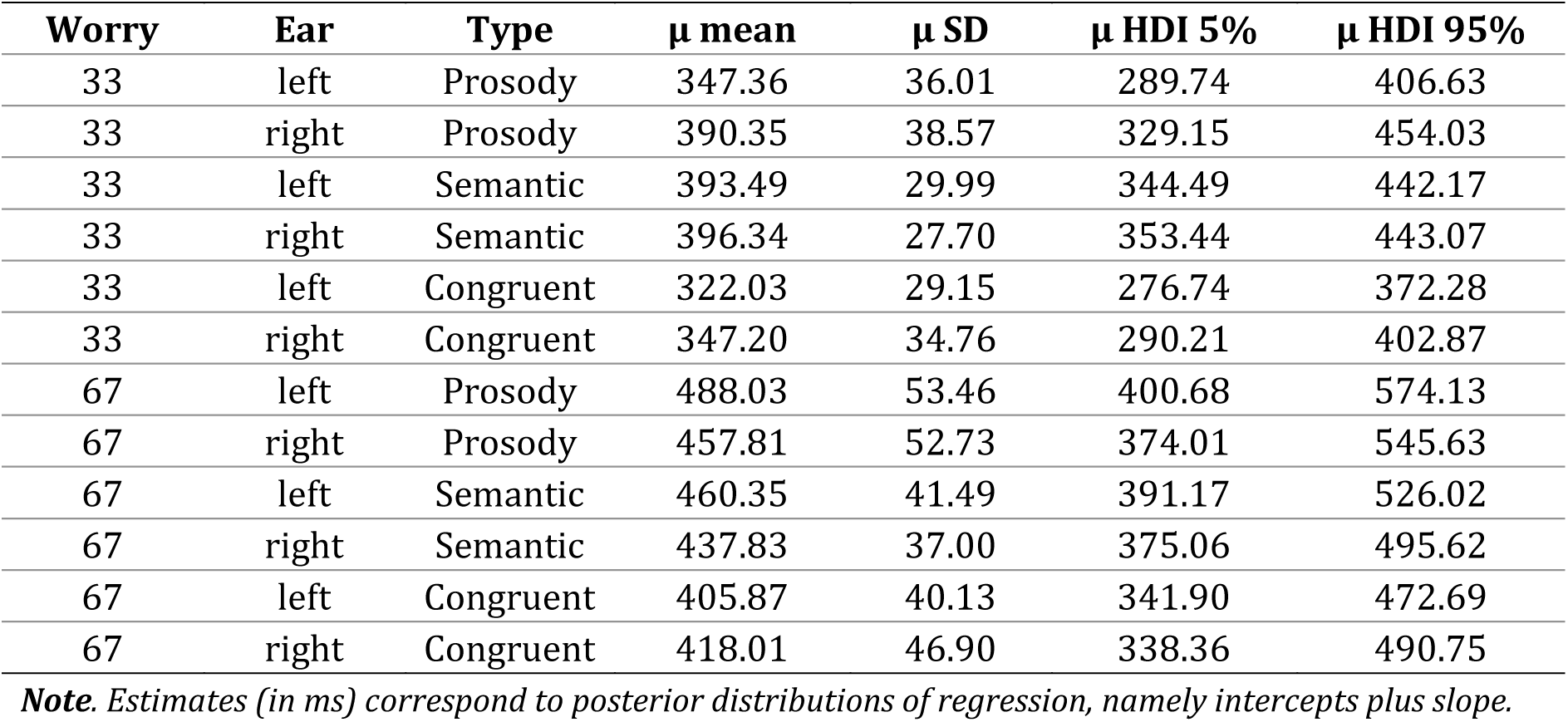
Experiment 1 (delayed response) indirect-threat reaction time estimates

| Worry | Ear | Type | $\mu$ mean | $\mu$ SD | $\mu$ HDI 5% | $\mu$ HDI 95% |
| --- | --- | --- | --- | --- | --- | --- |
| 33 | left | Prosody | 347.36 | 36.01 | 289.74 | 406.63 |
| 33 | right | Prosody | 390.35 | 38.57 | 329.15 | 454.03 |
| 33 | left | Semantic | 393.49 | 29.99 | 344.49 | 442.17 |
| 33 | right | Semantic | 396.34 | 27.70 | 353.44 | 443.07 |
| 33 | left | Congruent | 322.03 | 29.15 | 276.74 | 372.28 |
| 33 | right | Congruent | 347.20 | 34.76 | 290.21 | 402.87 |
| 67 | left | Prosody | 488.03 | 53.46 | 400.68 | 574.13 |
| 67 | right | Prosody | 457.81 | 52.73 | 374.01 | 545.63 |
| 67 | left | Semantic | 460.35 | 41.49 | 391.17 | 526.02 |
| 67 | right | Semantic | 437.83 | 37.00 | 375.06 | 495.62 |
| 67 | left | Congruent | 405.87 | 40.13 | 341.90 | 472.69 |
| 67 | right | Congruent | 418.01 | 46.90 | 338.36 | 490.75 |

**Note.** Estimates (in ms) correspond to posterior distributions of regression, namely intercepts plus slope.

Accuracy results for the direct-threat task indicate that responses to all conditions are estimated to be over 70% accurate and, excepting Semantic, they show a small but uncertain increase. Conditions at the right ear show to be around 10% less accurate respect to contralateral ear, but with substantial HDI overlap. See Table 4 and Figure 5 for summaries.

**Figure 5.**
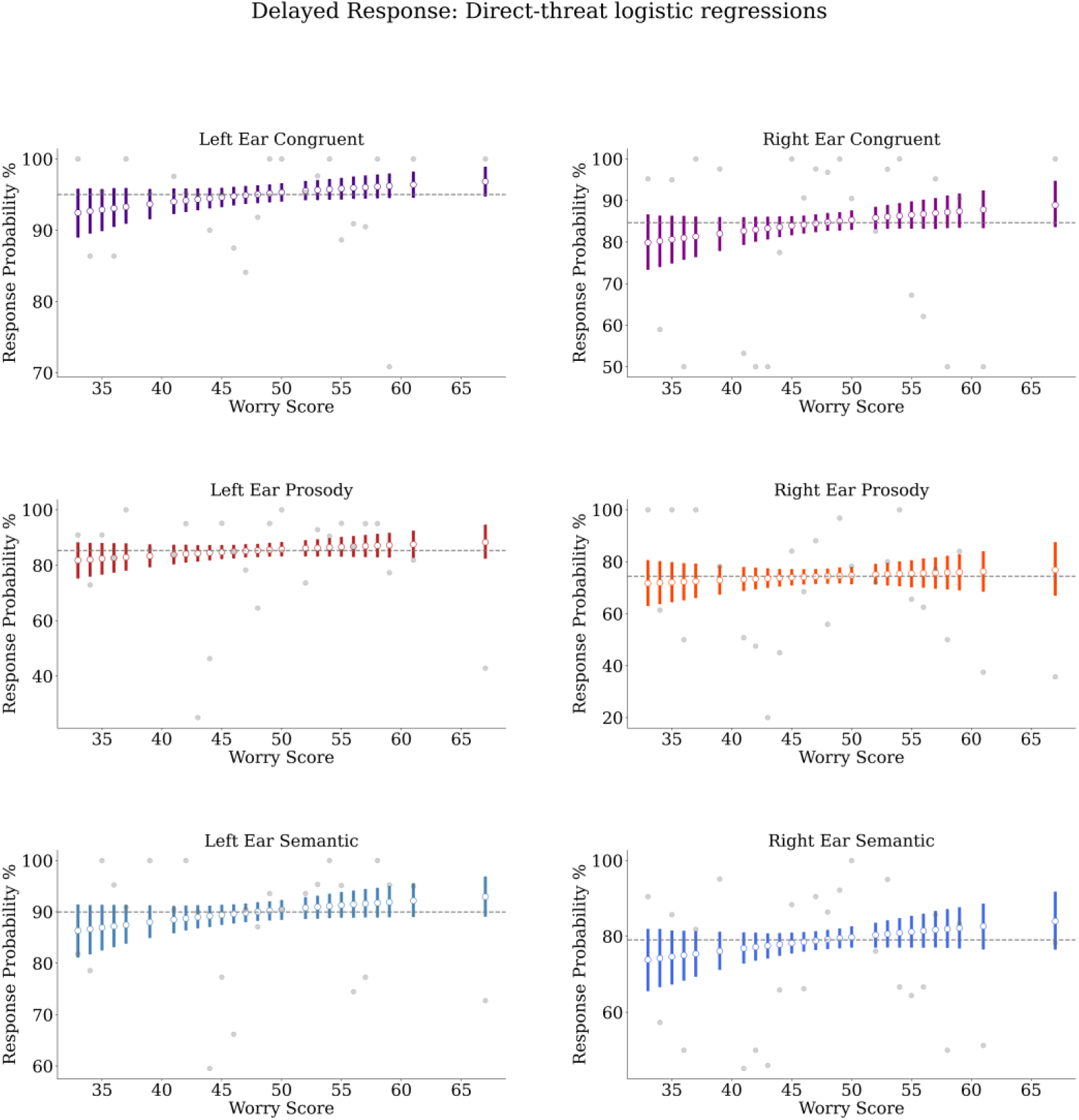
Experiment 1 (delayed response), direct-threat logistic regression estimates. White circles indicate posterior means. Coloured bars indicate highest density intervals (HDIs). Grey line indicates estimated worry score median’s posterior median. Grey dots indicate raw means. Note that estimates show a slight increase as a function of worry, but HDIs overlap in all conditions.

**Table 4.** Experiment 1 (delayed response) direct-threat accuracy estimates

| Worry | Ear | Type | $\mu$ mean | $\mu$ SD | $\mu$ HDI 5% | $\mu$ HDI 95% |
| --- | --- | --- | --- | --- | --- | --- |
| 33 | left | Prosody | 81.80 | 3.75 | 75.73 | 87.75 |
| 33 | right | Prosody | 71.73 | 5.04 | 63.52 | 80.07 |
| 33 | left | Semantic | 86.34 | 3.04 | 81.39 | 91.14 |
| 33 | right | Semantic | 73.83 | 4.77 | 65.93 | 81.59 |
| 33 | left | Congruent | 92.45 | 2.07 | 89.16 | 95.61 |
| 33 | right | Congruent | 79.88 | 3.95 | 73.70 | 86.32 |
| 67 | left | Prosody | 88.34 | 3.60 | 82.91 | 94.15 |
| 67 | right | Prosody | 76.87 | 6.06 | 67.55 | 87.01 |
| 67 | left | Semantic | 92.93 | 2.38 | 89.33 | 96.61 |
| 67 | right | Semantic | 83.99 | 4.58 | 76.88 | 91.39 |
| 67 | left | Congruent | 96.82 | 1.27 | 94.94 | 98.70 |
| 67 | right | Congruent | 88.92 | 3.37 | 83.97 | 94.38 |

**Note.** Estimates (in %) correspond to posterior distributions of regression, namely intercepts plus slope.

Similarly, accuracy results for the indirect-threat task indicate that responses to all conditions are estimated to be over 70% accurate and. In this case, right ear shows small and relatively certain increases for Congruent and Semantic at the right ear (around 5% to 8%), and Prosody tends to slightly decrease at the left ear (around 14%). All other conditions show little to no effect of worry or ear. Table 5 and Figure 6 summarise these results.

**Figure 6.**
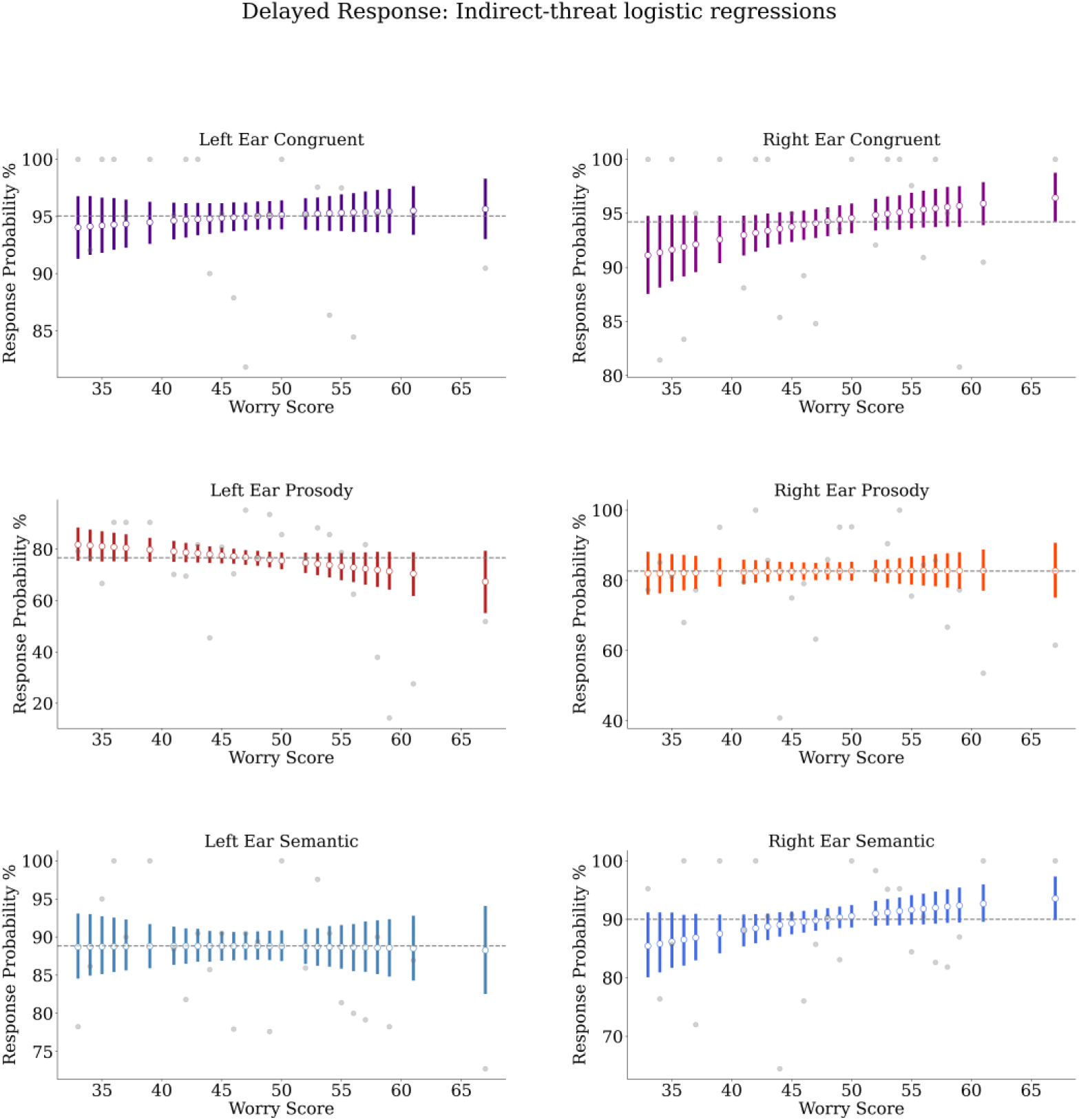
Experiment 1 (delayed response), indirect-threat logistic regression estimates. White circles indicate posterior means. Coloured bars indicate highest density intervals (HDIs). Grey line indicates estimated worry score median’s posterior median. Grey dots indicate raw means. Note that all conditions show slight increases or no increase, with general HDI overlap, excepting left ear Prosody that decreases (with some HDI overlap).

**Table 5.** Experiment 1 (delayed response) indirect-threat accuracy estimates

| **Worry** | **Ear** | **Type** | **$\mu$ mean** | **$\mu$ SD** | **$\mu$ HDI 5%** | **$\mu$ HDI 95%** |
| --- | --- | --- | --- | --- | --- | --- |
| 33 | left | Prosody | 81.81 | 3.70 | 76.10 | 87.89 |
| 33 | right | Prosody | 81.98 | 3.55 | 76.39 | 87.70 |
| 33 | left | Semantic | 88.70 | 2.55 | 84.74 | 92.89 |
| 33 | right | Semantic | 85.43 | 3.35 | 80.33 | 90.92 |
| 33 | left | Congruent | 94.05 | 1.66 | 91.43 | 96.64 |
| 33 | right | Congruent | 91.12 | 2.24 | 87.69 | 94.63 |
| 67 | left | Prosody | 67.36 | 7.07 | 55.69 | 78.85 |
| 67 | right | Prosody | 82.70 | 4.61 | 75.54 | 90.31 |
| 67 | left | Semantic | 88.26 | 3.57 | 82.74 | 93.88 |
| 67 | right | Semantic | 93.56 | 2.25 | 90.12 | 97.07 |
| 67 | left | Congruent | 95.65 | 1.67 | 93.15 | 98.18 |
| 67 | right | Congruent | 96.44 | 1.35 | 94.42 | 98.62 |

**Note.** Estimates (in %) correspond to posterior distributions of regression, namely intercepts plus slope.

### Experiment 2: Fast Response

Models for Experiment 2 did not include duration in the regression, as the relationship between duration and worry level is not clear, as participants must always answer before the end of each sentence. Again, all effects show good precision and all models sampled properly (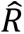 ≤ 1, ESS > 400), with energy plots, traceplots, and autocorrelation plots showing good convergence (for images see our OSF repository, link in the Data Statement section).

Results from the direct-threat task indicate that RTs consistently decrease as a function of worry for all ears and conditions. Congruent shows some HDI overlap, indicating less certain decreases. Instead, Prosody and Semantic decrease between 170ms and 213ms. Semantic and Congruent show faster RTs respect to Prosody, with differences ranging around 200ms to 300ms. Note that Prosody and Semantic, though showing wider HDIs at worry score extremes (less precision), show no HDI overlap, indicating better certainty for estimated decreases. See Table 6 and Figure 7 for summaries of these results.

**Figure 7.**
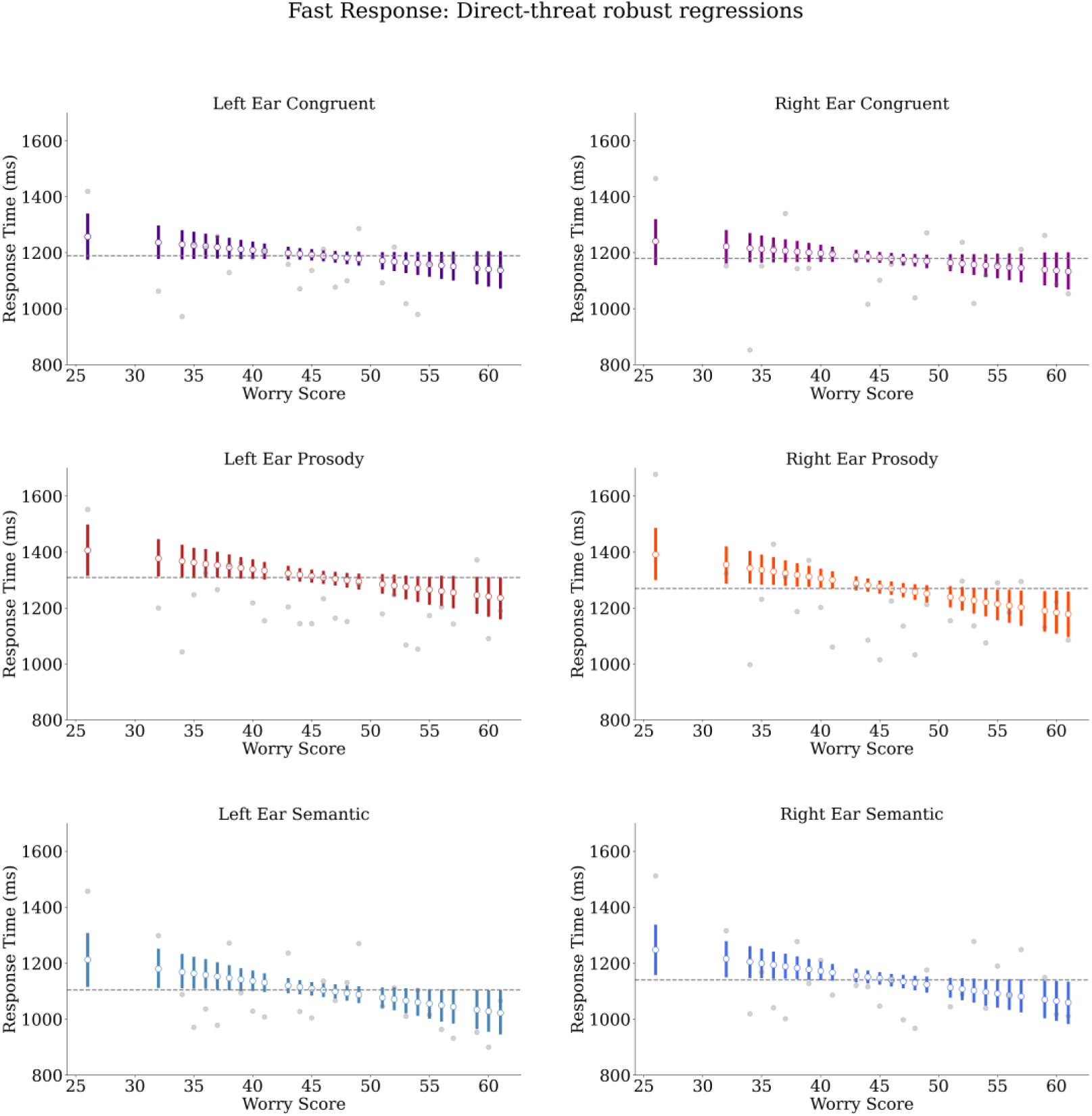
Experiment 2 (fast response), direct-threat robust regression estimates. White circles indicate posterior means. Coloured bars indicate highest density intervals (HDIs). Grey line indicates estimated worry score median’s posterior median. Grey dots indicate raw means. Note that estimates decrease as a function of worry, with little to no HDI overlap in all conditions, but Congruent shows a slightly smaller effect.

**Table 6.** Experiment 2 (fast response) direct-threat reaction time estimates

| Worry | Ear | Type | $\mu$ mean | $\mu$ SD | $\mu$ HDI 5% | $\mu$ HDI 95% |
| --- | --- | --- | --- | --- | --- | --- |
| 26 | left | Prosody | 1406.53 | 52.41 | 1320.33 | 1493.42 |
| 26 | right | Prosody | 1391.62 | 53.68 | 1305.41 | 1481.28 |
| 26 | left | Semantic | 1212.60 | 55.68 | 1120.54 | 1302.88 |
| 26 | right | Semantic | 1247.89 | 51.88 | 1162.93 | 1332.22 |
| 26 | left | Congruent | 1257.33 | 46.79 | 1180.63 | 1335.36 |
| 26 | right | Congruent | 1241.05 | 47.10 | 1160.82 | 1314.79 |
| 61 | left | Prosody | 1236.18 | 42.25 | 1164.46 | 1303.96 |
| 61 | right | Prosody | 1178.12 | 45.86 | 1101.87 | 1253.80 |
| 61 | left | Semantic | 1023.02 | 45.05 | 950.99 | 1098.92 |
| 61 | right | Semantic | 1059.78 | 43.20 | 988.14 | 1129.10 |
| 61 | left | Congruent | 1137.52 | 37.69 | 1076.98 | 1200.10 |
| 61 | right | Congruent | 1133.26 | 37.68 | 1073.40 | 1196.29 |

**Note.** Estimates (in ms) correspond to posterior distributions of regression, namely intercepts plus slope.

In the indirect-threat task, estimates show much smaller decreases in RT as a function of worry. Where HDI overlap is present in all conditions. Again, Congruent and Semantic show smaller RTs respect to prosody, in a similar range as previously observed. Table 7 and Figure 8 contain summaries of these results.

**Figure 8.**
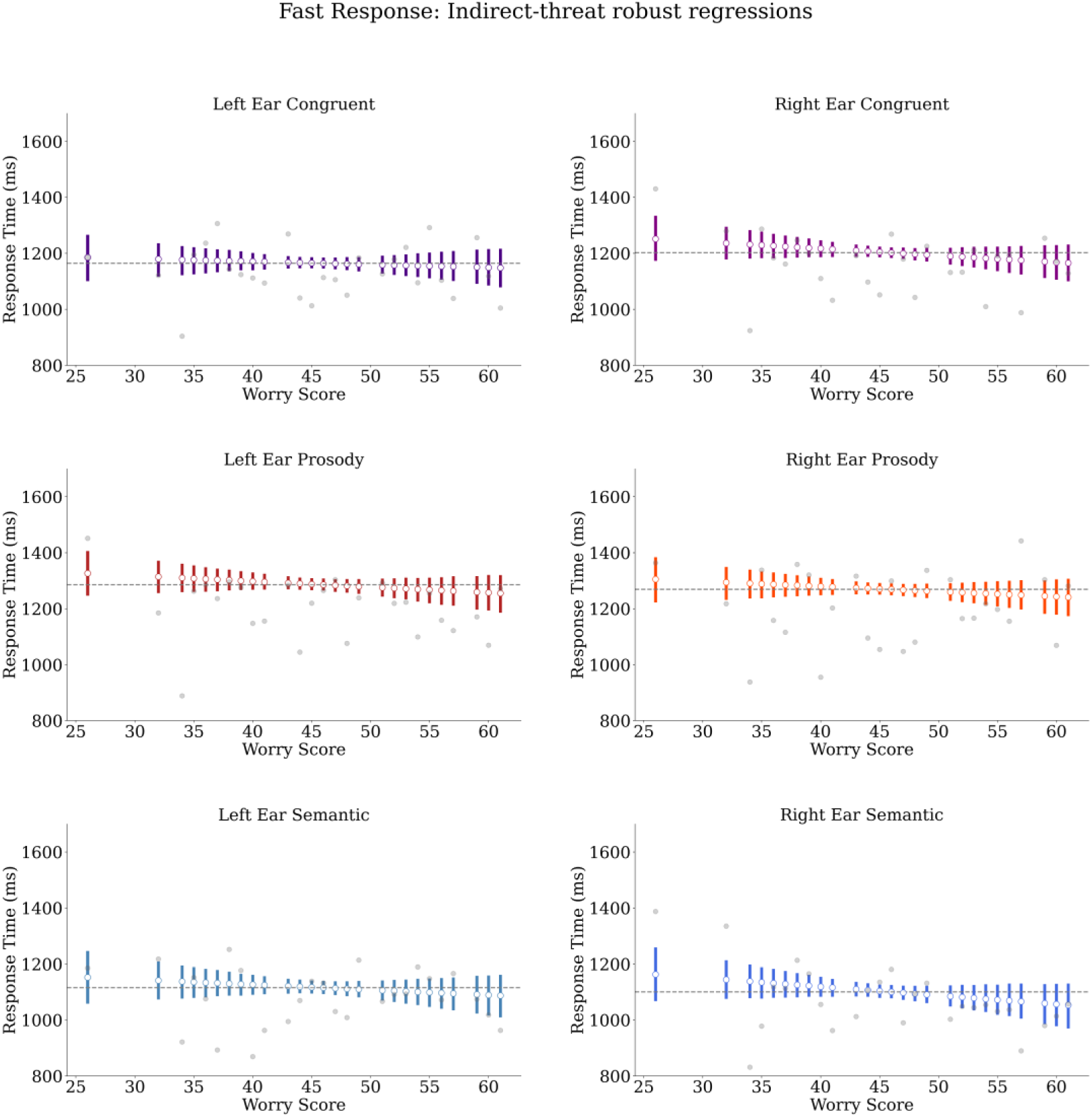
Experiment 2 (fast response), indirect-threat robust regression estimates. White circles indicate posterior means. Coloured bars indicate highest density intervals (HDIs). Grey line indicates estimated worry score median’s posterior median. Grey dots indicate raw means. Note that estimates decrease as a function of worry, but with substantial HDI overlap in all conditions.

**Figure 9.**
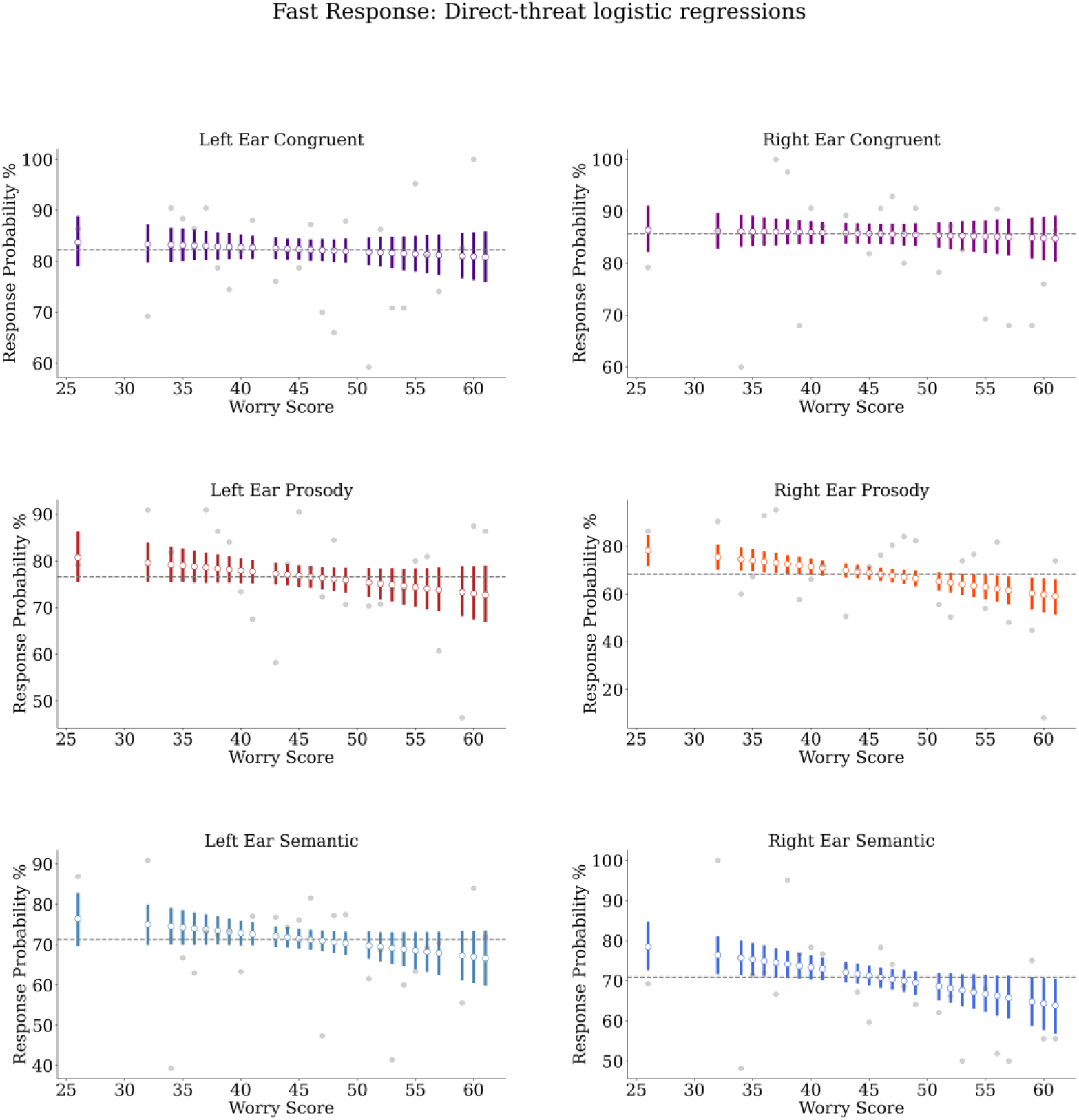
Experiment 2 (fast response), direct-threat logistic regression estimates. White circles indicate posterior means. Coloured bars indicate highest density intervals (HDIs). Grey line indicates estimated worry score median’s posterior median. Grey dots indicate raw means. Note that estimates show a slight decrease as a function of worry, but HDIs overlap in all conditions.

**Table 7.** Experiment 2 (fast response) indirect-threat reaction time estimates

| **Worry** | **Ear** | **Type** | **$\mu$ mean** | **$\mu$ SD** | **$\mu$ HDI 5%** | **$\mu$ HDI 95%** |
| --- | --- | --- | --- | --- | --- | --- |
| 26 | left | Prosody | 1326.40 | 46.01 | 1252.09 | 1401.28 |
| 26 | right | Prosody | 1305.27 | 46.56 | 1227.86 | 1378.76 |
| 26 | left | Semantic | 1152.51 | 54.75 | 1063.93 | 1240.96 |
| 26 | right | Semantic | 1163.11 | 55.56 | 1073.01 | 1253.78 |
| 26 | left | Congruent | 1185.62 | 47.73 | 1105.66 | 1261.18 |
| 26 | right | Congruent | 1251.36 | 46.15 | 1178.57 | 1329.32 |
| 61 | left | Prosody | 1255.32 | 37.90 | 1190.96 | 1314.64 |
| 61 | right | Prosody | 1241.50 | 37.71 | 1178.57 | 1301.70 |
| 61 | left | Semantic | 1087.65 | 44.17 | 1014.83 | 1156.30 |
| 61 | right | Semantic | 1053.64 | 46.23 | 975.16 | 1125.14 |
| 61 | left | Congruent | 1148.33 | 39.16 | 1083.64 | 1211.17 |
| 61 | right | Congruent | 1165.38 | 37.05 | 1105.18 | 1226.52 |

**Note.** Estimates (in ms) correspond to posterior distributions of regression, namely intercepts plus slope.

**Table 8.** Experiment 2 (fast response) direct-threat accuracy estimates

| Worry | Ear | Type | $\mu$ mean | $\mu$ SD | $\mu$ HDI 5% | $\mu$ HDI 95% |
| --- | --- | --- | --- | --- | --- | --- |
| 26 | left | Prosody | 80.78 | 3.18 | 75.73 | 85.99 |
| 26 | right | Prosody | 78.25 | 3.60 | 72.42 | 84.21 |
| 26 | left | Semantic | 76.45 | 3.89 | 70.02 | 82.51 |
| 26 | right | Semantic | 78.48 | 3.46 | 73.00 | 84.33 |
| 26 | left | Congruent | 83.76 | 2.89 | 79.30 | 88.55 |
| 26 | right | Congruent | 86.38 | 2.58 | 82.43 | 90.81 |
| 61 | left | Prosody | 72.77 | 3.46 | 67.32 | 78.69 |
| 61 | right | Prosody | 59.10 | 4.19 | 51.90 | 65.68 |
| 61 | left | Semantic | 66.60 | 3.94 | 60.17 | 73.11 |
| 61 | right | Semantic | 63.85 | 3.96 | 57.14 | 70.10 |
| 61 | left | Congruent | 80.83 | 2.88 | 76.24 | 85.57 |
| 61 | right | Congruent | 84.76 | 2.54 | 80.58 | 88.81 |

**Note.** Estimates (in %) correspond to posterior distributions of regression, namely intercepts plus slope.

Accuracy results for the direct-threat task indicate that responses to all conditions are estimated to be over 59% accurate. Congruent shows almost no change as a function of worry or ear. Prosody and Semantic at the left ear show decreases of around 8-9% respectively, but decreases of around 14% to 19% respectively at the right ear. Note that the latter show no HDI overlap. See Table 4 and Figure 5 for summaries.

Accuracy results for the indirect-threat condition also indicate overall 59% accuracy. All conditions seem to decrease as a function of worry (less than 10%), with Semantic at the right ear being the only condition without HDI overlap and showing a decrease of around 12% from lowest to highest worry levels. See Table 9 and Figure 10 for summaries of these results.

**Figure 10.**
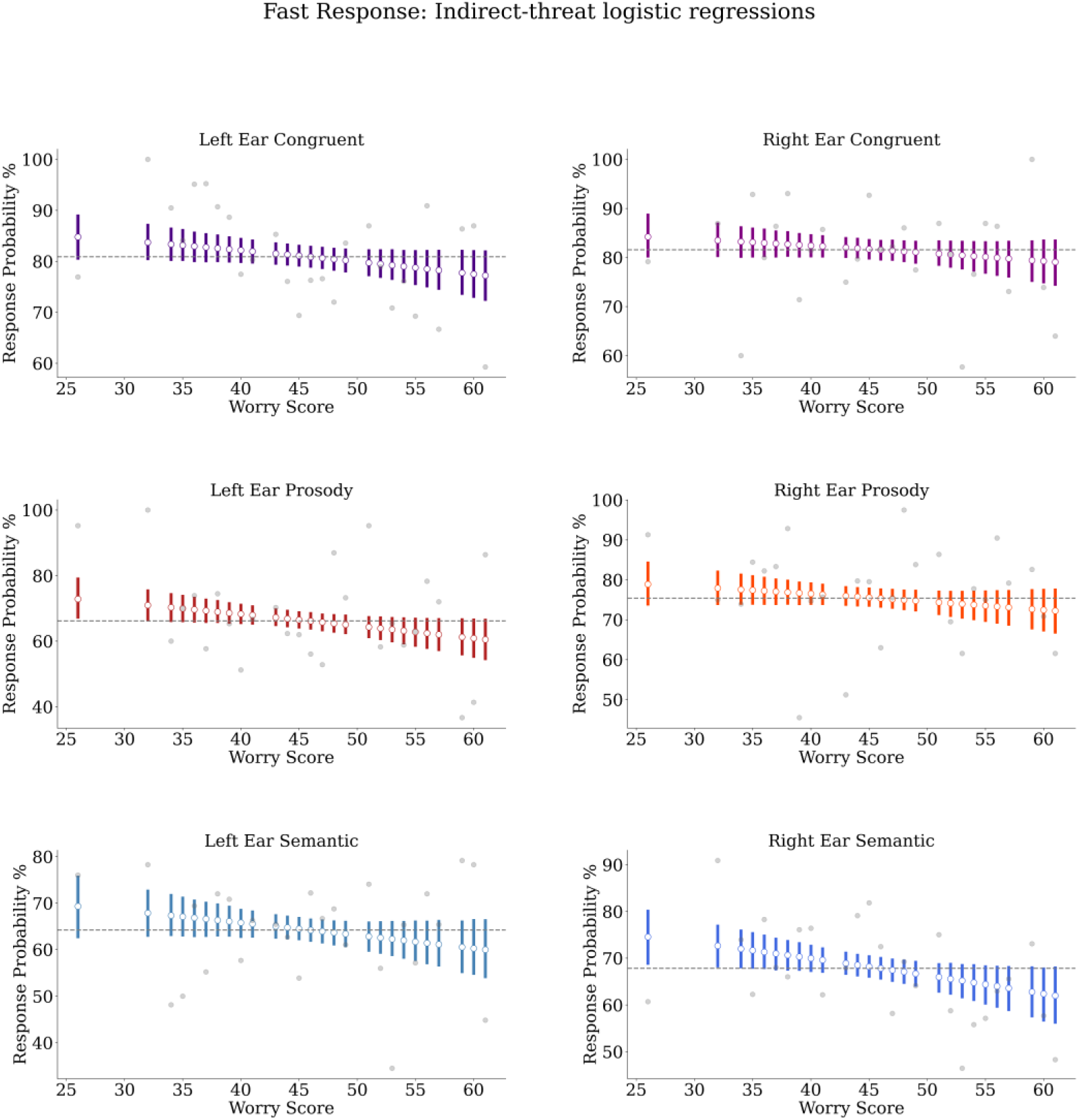
Experiment 2 (fast response), indirect-threat logistic regression estimates. White circles indicate posterior means. Coloured bars indicate highest density intervals (HDIs). Grey line indicates estimated worry score median’s posterior median. Grey dots indicate raw means. Note that estimates show a slight decrease as a function of worry, but HDIs overlap in most conditions, excepting left ear prosody and right ear semantic, which show little to no overlap.

**Table 9.** Experiment 2 (fast response) indirect-threat accuracy estimates

| **Worry** | **Ear** | **Type** | **$\mu$ mean** | **$\mu$ SD** | **$\mu$ HDI 5%** | **$\mu$ HDI 95%** |
| --- | --- | --- | --- | --- | --- | --- |
| 26 | left | Prosody | 72.84 | 3.52 | 67.32 | 78.96 |
| 26 | right | Prosody | 78.90 | 3.19 | 73.91 | 84.22 |
| 26 | left | Semantic | 69.30 | 3.92 | 62.77 | 75.58 |
| 26 | right | Semantic | 74.53 | 3.43 | 68.88 | 80.05 |
| 26 | left | Congruent | 84.78 | 2.54 | 80.62 | 88.91 |
| 26 | right | Congruent | 84.23 | 2.57 | 80.30 | 88.64 |
| 61 | left | Prosody | 60.48 | 3.55 | 54.68 | 66.43 |
| 61 | right | Prosody | 72.22 | 3.22 | 66.90 | 77.44 |
| 61 | left | Semantic | 59.97 | 3.63 | 54.16 | 66.22 |
| 61 | right | Semantic | 62.01 | 3.53 | 56.30 | 67.89 |
| 61 | left | Congruent | 77.22 | 2.83 | 72.52 | 81.87 |
| 61 | right | Congruent | 79.10 | 2.69 | 74.56 | 83.40 |

**Note.** Estimates (in %) correspond to posterior distributions of regression, namely intercepts plus slope.

## Discussion

Results from the delayed response experiment (Experiment 1) indicate small effects of worry on RT and accuracy. In the direct-threat task RTs tend to increase slightly more at the right ear as a function of worry, in particular for Prosody. In the indirect-threat task, RTs increase generally more as a function of worry, in particular for the left ear and the Prosody condition. However, for both tasks, effects are considerably small as evidenced by relatively high uncertainty and HDI overlap. Accuracy results from direct-threat indicate small increases as a function of worry for all conditions, with slight but uncertain underperformance at the right ear for all sentence types. Results from indirect-threat indicate less certainty and a less coherent pattern, which may be indicative of no systematic effect. Results from the fast response experiment (Experiment 2) showed the opposite pattern. Direct-threat results indicate that RTs become faster as a function of worry. These effects are generally certain for Prosody and Semantic types at both ears. At the indirect-threat task, results indicate that effects are considerable reduced for all conditions, namely RTs decrease less. Direct-threat accuracy tends to decrease as a function of worry, with more certain effects for right ear Prosody and Semantic. Indirect-threat results indicate a reduction of effects and certainty, but only for left ear Prosody and right ear Semantic.

For present purposes we will take the most conservative approach. That is, we uphold the most straightforward interpretation, which takes all effects into account simultaneously due to their involvement into an interaction. This simply indicates that Experiment 1 (delayed response) results need to be considered as generally uncertain, and only Experiment 2 (fast response) direct-threat results can be considered relevant. In the latter task, Congruent RT results do not show very certain effects, but RTs of Prosody and Semantic at both ears tend to decrease up to around 200ms as a function of worry (PSWQ score), Prosody RTs are 100ms greater overall respect to other conditions (though with some HDI overlap). For the same experiment and task, accuracy shows to relevantly decrease only for Prosody and Semantics at the right ear. This implies that our hypotheses indicating ear preferences as distinct by type cannot be supported. However, our hypothesis on differing effects of fast responses is partially corroborated. This, however, is linked to an effect of congruency rather than emotional information type. In other words, when threatening prosody and semantics match (Congruent type), effects of worry are minimal, and when they do not match (Prosody and Semantic), worry induces faster responses (smaller RTs). Furthermore, this can evidence an accuracy-speed trade-off as faster responses tend to be associated with decreased accuracy, but only at the right ear.

In order put the present results in context, it is important to recapitulate important aspects that differentiate the current experiments from previous relevant studies: 1) The use of worry-level as a continuous variable. Worry is associated with anxious apprehension (Heller et al, 1997), which implies more chances of participants over-engaging with threat. 2) Stimuli were semi-naturalistic sentences, providing stronger contextual effects. In addition, their longer durations can facilitate engagement with their content. 3) Information channels were manipulated to disentangle effects of semantics and prosody from effects of emotional expression (Kotz and Paulmann, 2007). 4) The use of two tasks measuring responses directed to threatening or neutral stimuli (direct vs. indirect threat, e.g. Sanders et al, 2005) helps to check whether attention effects could be inducing different response patterns. 5) Two experiments were implemented to verify whether answering after sentences’ end or during sentence presentations (delayed vs. fast) can influence laterality patterns by tapping into different moments of a multistep emotional language processing mechanism (Kotz and Paulmann, 2011).

With this in mind, it is important to carefully interpret the lack of clear laterality (ear) effects for most tasks. Weak ear effects might be explained by the great variability between items and the high duration (also very variable) of sentences. However, the lack of sensitivity of DL when more naturalistic stimuli/context are provided cannot be discarded as a possible explanation. If DL effects are task dependent (Godfrey and Grimshaw, 2015), increased naturalness on stimuli and context can bring out a myriad of bilateral processing patterns that might make ear advantages disappear on the long run when prolonged auditory stimuli are listened to. Although there is previous evidence suggesting a right lateralized pattern for prosody vs. semantic evaluation in an EEG experiment (not considering anxiety), using a congruency (not DL) task with sentences as stimuli (Kotz and Paulmann, 2007), further experimentation using a similar paradigm has not observed this pattern (Paulmann and Kotz, 2012). Although this pattern is explained by the strong association between pitch recognition and RH engagement (Kotz and Paulmann, 2007; Zatorre et al., 2002), there are other frequency and spectral features that might be important for recognizing both threatening and neutral sentences (Banse and Scherer, 1996; Hammerschmidt and Jürgens, 2007; Liu et al., 2013; Xu et al., 2013; Zatorre et al., 2002). This could imply that distinguishing prosody and semantics might be a continuous process that can have diversified effects even during sentence presentation.

Indeed, by manipulating angry prosody changes at the beginning and end of sentences, an EEG study has observed that when prosody changes from angry to neutral within sentences, processing is more effortful (Chen et al, 2011). This might indicate that the rich acoustic nature of prosody might be detected quickly but resourcefully. Recent EEG research has observed that anxious people present ERP differences at both early and late processing stages when answering to threatening prosody and non-language vocalizations (Pell et al, 2015). This is consistent with the notion of early over-attention and later over-engagement, and indicates that behavioural responses might change given early or late variations in threat.

Another possible explanation is callosal relay (Atchely et al., 2011; Grimshaw et al, 2003), where increased anxiety would disrupt RH to LH callosal information transferring of threatening prosody. It has been proposed that callosal relay is highly relevant for language informational and emotional processing (Friederici et al, 2007; Kotz and Paulmann, 2011; Steinmann et al., 2017). Hence, interference at one hemisphere (e.g. rumination or worry impacting LH) can have an effect on information transferring to the other. Thus, callosal relay effects could have a relevant impact on how DL tasks are processed, subject to both top-down and bottom-up effects (Westernhausen and Hugdahl, 2008), which is particularly relevant when laterality effects induced by acoustic or lexical properties need to be disentangled from those induced solely by emotional processing (Grimshaw et al, 2003; Leshem, 2018).

These patterns indicate that our extension of dichotic listening models (Grimshaw et al., 2003) does not guarantee laterality effects induced by sentences’ information type. Also, our prediction of a strong effect of worry level (trait anxiety), affecting emotional language processing due to possible over-engagement with threat (Bar-Haim et al., 2007; Spielberg et al., 2013), was not strongly supported. It was proposed that delayed responses facilitate over-engagement with threat due to the long latency between sentence presentation and response. This, together with the high variability in sentences’ durations and content might have nullified ear and/or type effects and may not be enough to induce a sufficiently strong over-engagement-related delay in responses.

Although there is a trend indicating a response slow-down as a function of worry for direct-threat right ear Prosody and indirect-threat left ear prosody, more certain results are required to confirm these patterns. This implies that this trend may not be replicated in future studies, thus we do not develop further interpretation. Furthermore, in Experiment 2, we failed to observe any clear Type or ear effect when responses were forced to be fast (during sentence), besides a small accuracy effect on ear at the direct-threat task for Semantic and Prosody only. Partially aligned with our prediction, the pattern of Experiment 2 (fast response) is the opposite as in Experiment 1 (delayed response). That is, RTs decrease as a function of worry (apprehensive anxiety) and accuracy also decreases. This, however, may evidence an accuracy-speed trade-off related with a general effect task effect (Robinson et al., 2013), where more anxious participants tend to sacrifice accuracy to answer faster to stimuli presented under pressure. Thus, it is difficult to corroborate our hypothesis about specific early or mid-early effects of emotional speech. First, RT increases are associated with information type (semantic or prosodic), but with congruency. Second, the effects happen irrespective of ear, giving no evidence of lateralisation effects. Even so accuracy tends to show a more consistent effect at the right ear only, this does not match a hypothetical RH privilege for either prosodic or emotional content (Kotz and Paulmann, 2011).

Similarly, previous research using single words, dichotically presented as direct- and indirect-threat (or anger), and measuring anxiety, did not find differences in RT for left or right ears (Sander et al., 2005; Leshem, 2018; Peshard, 2016), but did find differences in attention focus per ear. Present results do not indicate a clear distinct pattern on indirect-threat tasks. Recent research (Leshem, 2018) did not find effects of trait anxiety on ear either; present results, going even further, evidence a precise pattern of weak or negligible interactions between ear and worry (trait anxiety). Although, it is also important to emphasize that the absence of other effects might be induced by stimuli’s high variability in length and content. This lack of clear ear effects makes difficult to integrate DL literature to explain whether this RT/accuracy decrease is actually associated with effects of anxiety on early-mid speech processing stages of language processing, as we proposed based on multistep models of emotional language (Kotz and Paulmann, 2011).

Given this, our results suggest that any type of threatening language, attended either directly or indirectly, is affected by apprehensive anxiety only under task pressure (i.e. fast response). Therefore, our proposal of adding a fourth stage to a multistep model of emotional language (Kotz and Paulmann, 2011) is not well supported. Previous research has found that non-language simple threatening stimuli (e.g. noise) can indeed induce similar over-engagement effects (slower RTs) in association with behavioural inhibition but not with trait anxiety (Massar et al., 2011), while other studies indicate that induced anxiety slows down RTs irrespective of stimulus emotional content (Aylward et al., 2017). Presently, by using longer and more naturalistic stimuli, we have observed the opposite pattern. Only direct attention (answering directly) to threat induces a speed-up in RTs and a decrease in accuracy when fast responses are required and stimuli are not congruent (both semantically and prosodically threatening). Note that present congruency is informational (prosodic-semantic) not emotional congruency as explored in previous research on induced anxiety (e.g. Robinson et al., 2012). This may imply that attention mechanism of early detection, namely over-attention to threat, could provide an explanation for faster responses when the stimulation period is shortened or responses are under pressure (Cisler and Koster, 2010), in particular for stimuli which are harder to recognise/process (i.e. not congruent).

Therefore, our initial assumption that a fast response experiment (Experiment 2) would be enough to identify difference at early processing stages is not necessarily correct, at least given present stimuli and task. The varied position of threatening lexical items and/or threatening intonation emphasis might cause a general slow-down of responses, because very specific features of sentences need to be identified and participants have time to do so (the whole extent of a sentence). While evaluation mechanisms could serve as an explanation, the fact that there are not strong effects associated with difficulties categorising of identifying stimuli makes them weak candidates. This may also be related to a general caveat of our approach. Behavioural measures such as DL, though able to portray a very general picture of underlying brain processes, might not be enough. Better spatial and temporal resolution is required to disentangle laterality and early-stage effects of threatening language. The latter is particularly relevant, as the time-course of emotional language processing might have crucial differences at much shorter time-scales, as evidenced by previous EEG research (Chen et al, 2011; Kotz and Paulmann, 2007; Paulmann and Kotz, 2012; Pell et al, 2015; Wabnitz et al., 2015; Wambacq and Jerger, 2004). In consequence, present tasks could be replicated by using EEG measures, in particular Experiment 1, where EEG measures such as event-related potentials could provide richer information about processing occurring during sentence listening, before response preparation and response execution. This could also provide lab results as point of comparison with present web-based results. But more importantly, this is crucial for identifying differences in the neural signature of anxiety and language processing, indispensable for properly understanding time-related models of language and anxiety processing.

In conclusion, present results indicate that extending multistep models of language processing (Schirmer and Kotz, 2006; Kotz and Paulmann, 2011) by including aspects of multistage models of anxiety (Bar-Haim et al, 2007; Corr and McNaughton, 2012) could be a relevant theoretical approach. However, present results do not provide strong support for present hypotheses, which makes necessary to test the language-anxiety relationship in different ways. Present studies suggest a speed-accuracy trade-off as a function of anxiety when responses are required to be fast, which mainly affects stimuli lacking one informational dimension (i.e. prosody-only threat or semantic-only threat). Nevertheless, these are not sufficient to ascertain a relationship between language and anxiety processing. Further experimental testing is thus required, in particular by implementing physiological measures such as EEG, and tasks that do not involve DL, using more controlled stimuli and investigating the effects of stimuli below or above the sentence level, such as phrases or narratives.

## Data statement

All data, analyses’ scripts, and additional info can be found at the Open Science Framework (OSF) repository: https://osf.io/z8pgf/?view_only=b5da5ce6c8644bc182231cd9b96be173. Identifier: DOI 10.17605/OSF.IO/Z8PGF.

## Acknowledgements

This research is supported by CONICYT Becas Chile 72170145 (https://www.conicyt.cl/). Contributions to the study were as follows. **Busch-Moreno**: Conceptualization, Methodology, Software, Analysis, Investigation, Resources, Curation, Writing, Visualization, Administration. **Vinson**: Conceptualization, Methodology, Resources, Revision, Supervision, Administration. **Tuomainen**: Conceptualization, Resources, Revision, Supervision, Administration. Preliminary results of this experiment, using a frequentist framework, were presented at the International Congress of Phonetic Sciences (ICPhS), Melbourne, 2019.

## References

Atchley, R. A., Grimshaw, G., Schuster, J., & Gibson, L. (2011). Examining lateralized lexical ambiguity processing using dichotic and cross-modal tasks. Neuropsychologia, 49(5), 1044–1051. https://doi.org/10.1016/j.neuropsychologia.2011.01.010

Aue, T., Cuny, C., Sander, D., & Grandjean, D. (2011). Peripheral responses to attended and unattended angry prosody: A dichotic listening paradigm. Psychophysiology, 48(3), 385–392. https://doi.org/10.1111/j.1469-8986.2010.01064.x

Aylward, J., Valton, V., Goer, F., Mkrtchian, A., Lally, N., Peters, S., Limbachya, T., & Robinson, O. J. (2017). The impact of induced anxiety on affective response inhibition. Royal Society Open Science, 4(6), 170084. https://doi.org/10.1098/rsos.170084

Banse, R., & Scherer, K. R. (1996). Acoustic profiles in vocal emotion expression. Journal of Personality and Social Psychology, 70(3), 614–636. https://doi.org/10.1037//0022-3514.70.3.614

Bar-Haim, Y., Lamy, D., Pergamin, L., Bakermans-Kranenburg, M. J., & van IJzendoorn, M. H. (2007). Threat-related attentional bias in anxious and nonanxious individuals: A meta-analytic study. Psychological Bulletin, 133(1), 1–24. https://doi.org/10.1037/0033-2909.133.1.1

Bates, D., Mächler, M., Bolker, B., & Walker, S. (2015). Fitting Linear Mixed-Effects Models Usinglme4. Journal of Statistical Software, 67(1). https://doi.org/10.18637/jss.v067.i01

Belin, P., Fecteau, S., & Bédard, C. (2004). Thinking the voice: neural correlates of voice perception. Trends in Cognitive Sciences, 8(3), 129–135. https://doi.org/10.1016/j.tics.2004.01.008

Borelli, E., Crepaldi, D., Porro, C. A., & Cacciari, C. (2018). The psycholinguistic and affective structure of words conveying pain. PLOS ONE, 13(6), e0199658. https://doi.org/10.1371/journal.pone.0199658

Bruder, G. E., Wexler, B. E., Stewart, J. W., Price, L. H., & Quitkin, F. M. (1999). Perceptual asymmetry differences between major depression with or without a comorbid anxiety disorder: A dichotic listening study. Journal of Abnormal Psychology, 108(2), 233–239. https://doi.org/10.1037/0021-843x.108.2.233

Chen, X., Zhao, L., Jiang, A., & Yang, Y. (2011). Event-related potential correlates of the expectancy violation effect during emotional prosody processing. Biological Psychology, 86(3), 158–167. https://doi.org/10.1016/j.biopsycho.2010.11.004

Corr, P. J., & McNaughton, N. (2012). Neuroscience and approach/avoidance personality traits: A two stage (valuation–motivation) approach. Neuroscience & Biobehavioral Reviews, 36(10), 2339–2354. https://doi.org/10.1016/j.neubiorev.2012.09.013

Davidson, R. J. (1992). Emotion and Affective Style: Hemispheric Substrates. Psychological Science, 3(1), 39–43. https://doi.org/10.1111/j.1467-9280.1992.tb00254.x

Eldar, S., Yankelevitch, R., Lamy, D., & Bar-Haim, Y. (2010). Enhanced neural reactivity and selective attention to threat in anxiety. Biological Psychology, 85(2), 252–257. https://doi.org/10.1016/j.biopsycho.2010.07.010

Engels, A. S., Heller, W., Mohanty, A., Herrington, J. D., Banich, M. T., Webb, A. G., & Miller, G. A. (2007). Specificity of regional brain activity in anxiety types during emotion processing. Psychophysiology, 44(3), 352–363. https://doi.org/10.1111/j.1469-8986.2007.00518.x

Friederici, A. D., von Cramon, D. Y., & Kotz, S. A. (2007). Role of the Corpus Callosum in Speech Comprehension: Interfacing Syntax and Prosody. Neuron, 53(1), 135–145. https://doi.org/10.1016/j.neuron.2006.11.020

Gadea, M., Espert, R., Salvador, A., & Martí-Bonmatí, L. (2011). The sad, the angry, and the asymmetrical brain: Dichotic Listening studies of negative affect and depression. Brain and Cognition, 76(2), 294–299. https://doi.org/10.1016/j.bandc.2011.03.003

Godfrey, H. K., & Grimshaw, G. M. (2015). Emotional language is all right: Emotional prosody reduces hemispheric asymmetry for linguistic processing. Laterality: Asymmetries of Body, Brain and Cognition, 21(4–6), 568–584. https://doi.org/10.1080/1357650x.2015.1096940

Grimshaw, G. M., Kwasny, K. M., Covell, E., & Johnson, R. A. (2003). The dynamic nature of language lateralization: effects of lexical and prosodic factors. Neuropsychologia, 41(8), 1008–1019. https://doi.org/10.1016/s0028-3932(02)00315-9

Grimshaw, G. M., Séguin, J. A., & Godfrey, H. K. (2009). Once more with feeling: The effects of emotional prosody on hemispheric specialisation for linguistic processing. Journal of Neurolinguistics, 22(4), 313–326. https://doi.org/10.1016/j.jneuroling.2008.10.005

Hammerschmidt, K., & Jürgens, U. (2007). Acoustical Correlates of Affective Prosody. Journal of Voice, 21(5), 531–540. https://doi.org/10.1016/j.jvoice.2006.03.002

Heller, W., Nitschke, J. B., Etienne, M. A., & Miller, G. A. (1997). Patterns of regional brain activity differentiate types of anxiety. Journal of Abnormal Psychology, 106(3), 376–385. https://doi.org/10.1037/0021-843x.106.3.376

Ho, S. M. Y., Mak, C. W. Y., Yeung, D., Duan, W., Tang, S., Yeung, J. C., & Ching, R. (2015). Emotional Valence, Arousal, and Threat Ratings of 160 Chinese Words among Adolescents. PLOS ONE, 10(7), e0132294. https://doi.org/10.1371/journal.pone.0132294

Hugdahl, K. (2011). Fifty years of dichotic listening research – Still going and going and…. Brain and Cognition, 76(2), 211–213. https://doi.org/10.1016/j.bandc.2011.03.006

Hunter, J. D. (2007). Matplotlib: A 2D Graphics Environment. Computing in Science & Engineering, 9(3), 90–95. https://doi.org/10.1109/mcse.2007.55

Jadoul, Y., Thompson, B., & de Boer, B. (2018). Introducing Parselmouth: A Python interface to Praat. Journal of Phonetics, 71, 1–15. https://doi.org/10.1016/j.wocn.2018.07.001

Kotz, S. A., & Paulmann, S. (2007). When emotional prosody and semantics dance cheek to cheek: ERP evidence. Brain Research, 1151, 107–118. https://doi.org/10.1016/j.brainres.2007.03.015

Kotz, S. A., & Paulmann, S. (2011). Emotion, Language, and the Brain. Language and Linguistics Compass, 5(3), 108–125. https://doi.org/10.1111/j.1749-818x.2010.00267.x

Kruschke, J. K. (2013). Bayesian estimation supersedes the t test. Journal of Experimental Psychology: General, 142(2), 573–603. https://doi.org/10.1037/a0029146

Kruschke, J. K. (2015). Doing Bayesian data analysis : a tutorial with R, JAGS, and stan. Amsterdam Etc: Elsevier, Academic Press, Cop.

Kruschke, J.K. (2018). Rejecting or Accepting Parameter Values in Bayesian Estimation. Advances in Methods and Practices in Psychological Science, 1(2), pp.270–280.

Kumar, R., Carroll, C., Hartikainen, A., & Martin, O. (2019). ArviZ a unified library for exploratory analysis of Bayesian models in Python. Journal of Open Source Software, 4(33), 1143. https://doi.org/10.21105/joss.01143

Leshem, R. (2018). Trait Anxiety and Attention: Cognitive Functioning as a Function of Attentional Demands. Current Psychology. https://doi.org/10.1007/s12144-018-9884-9

Liebenthal, E., Binder, J. R., Spitzer, S. M., Possing, E. T., & Medler, D. A. (2005). Neural Substrates of Phonemic Perception. Cerebral Cortex, 15(10), 1621–1631. https://doi.org/10.1093/cercor/bhi040

Liu, F., Xu, Y., Prom-on, S., & Yu, A. (2013). Morpheme-like prosodic functions: Evidence from acoustic analysis and computational modeling - UCL Discovery. Ucl.Ac.Uk. https://doi.org/http://discovery.ucl.ac.uk/1391681/1/Xu_Liu_etAl_JoSS2013.pdf

MacNamara, A., & Hajcak, G. (2010). Distinct electrocortical and behavioral evidence for increased attention to threat in generalized anxiety disorder. Depression and Anxiety, 27(3), 234–243. https://doi.org/10.1002/da.20679

Martin, O. (2018). Bayesian analysis with Python : introduction to statistical modeling and probabilistic programming using PyMC3 and ArviZ. Birmingham, Uk: Packt Publishing.

Massar, S. A. A., Mol, N. M., Kenemans, J. L., & Baas, J. M. P. (2011). Attentional bias in high- and low-anxious individuals: Evidence for threat-induced effects on engagement and disengagement. Cognition & Emotion, 25(5), 805–817. https://doi.org/10.1080/02699931.2010.515065

McElreath, R. (2020). Statistical rethinking : a Bayesian course with examples in R and Stan. Second Edition. Boca Raton ; London ; New York: Chapman & Hall/Crc.

McEvoy, P. M., Mahoney, A. E. J., & Moulds, M. L. (2010). Are worry, rumination, and post-event processing one and the same? Journal of Anxiety Disorders, 24(5), 509–519. https://doi.org/10.1016/j.janxdis.2010.03.008

McLaughlin, K. A., Borkovec, T. D., & Sibrava, N. J. (2007). The Effects of Worry and Rumination on Affect States and Cognitive Activity. Behavior Therapy, 38(1), 23–38. https://doi.org/10.1016/j.beth.2006.03.003

McNaughton, N., & Corr, P. J. (2014). Approach, avoidance, and their conflict: the problem of anchoring. Frontiers in Systems Neuroscience, 8. https://doi.org/10.3389/fnsys.2014.00124

McNaughton, N., & Gray, J. A. (2000). Anxiolytic action on the behavioural inhibition system implies multiple types of arousal contribute to anxiety. Journal of Affective Disorders, 61(3), 161–176. https://doi.org/10.1016/s0165-0327(00)00344-x

Meyer, T. J., Miller, M. L., Metzger, R. L., & Borkovec, T. D. (1990). Development and validation of the Penn State Worry Questionnaire. Behaviour Research and Therapy, 28(6), 487–495. Retrieved from https://www.ncbi.nlm.nih.gov/pubmed/2076086

Nitschke, J. B., Heller, W., Palmieri, P. A., & Miller, G. A. (1999). Contrasting patterns of brain activity in anxious apprehension and anxious arousal. Psychophysiology, 36(5), 628–637. https://doi.org/10.1111/1469-8986.3650628

Nygaard, L. C., Herold, D. S., & Namy, L. L. (2009). The Semantics of Prosody: Acoustic and Perceptual Evidence of Prosodic Correlates to Word Meaning. Cognitive Science, 33(1), 127–146. https://doi.org/10.1111/j.1551-6709.2008.01007.x

Paulmann, S., Jessen, S., & Kotz, S. A. (2012). It’s special the way you say it: An ERP investigation on the temporal dynamics of two types of prosody. Neuropsychologia, 50(7), 1609–1620. https://doi.org/10.1016/j.neuropsychologia.2012.03.014

Pell, M. D., Rothermich, K., Liu, P., Paulmann, S., Sethi, S., & Rigoulot, S. (2015). Preferential decoding of emotion from human non-linguistic vocalizations versus speech prosody. Biological Psychology, 111, 14–25. https://doi.org/10.1016/j.biopsycho.2015.08.008

Peschard, V., Gilboa-Schechtman, E., & Philippot, P. (2017). Selective attention to emotional prosody in social anxiety: a dichotic listening study. Cognition & Emotion, 31(8), 1749–1756. https://doi.org/10.1080/02699931.2016.1261012

Poeppel, D. (2003). The analysis of speech in different temporal integration windows: cerebral lateralization as ‘asymmetric sampling in time.’ Speech Communication, 41(1), 245–255. https://doi.org/10.1016/s0167-6393(02)00107-3

Poeppel, D., Idsardi, W. J., & van Wassenhove, V. (2008). Speech perception at the interface of neurobiology and linguistics. Philosophical Transactions of the Royal Society B: Biological Sciences, 363(1493), 1071–1086. https://doi.org/10.1098/rstb.2007.2160

Robinson, O. J., Krimsky, M., & Grillon, C. (2013). The impact of induced anxiety on response inhibition. Frontiers in Human Neuroscience, 7. https://doi.org/10.3389/fnhum.2013.00069

Ross, E. D., Thompson, R. D., & Yenkosky, J. (1997). Lateralization of Affective Prosody in Brain and the Callosal Integration of Hemispheric Language Functions. Brain and Language, 56(1), 27–54. https://doi.org/10.1006/brln.1997.1731

Salvatier, J., Wiecki, T. V., & Fonnesbeck, C. (2016). Probabilistic programming in Python using PyMC3. PeerJ Computer Science, 2, e55. https://doi.org/10.7717/peerj-cs.55

Sander, D., Grandjean, D., Pourtois, G., Schwartz, S., Seghier, M. L., Scherer, K. R., & Vuilleumier, P. (2005). Emotion and attention interactions in social cognition: Brain regions involved in processing anger prosody. NeuroImage, 28(4), 848–858. https://doi.org/10.1016/j.neuroimage.2005.06.023

Schirmer, A., & Kotz, S. A. (2003). ERP Evidence for a Sex-Specific Stroop Effect in Emotional Speech. Journal of Cognitive Neuroscience, 15(8), 1135–1148. https://doi.org/10.1162/089892903322598102

Schirmer, A., & Kotz, S. A. (2006). Beyond the right hemisphere: brain mechanisms mediating vocal emotional processing. Trends in Cognitive Sciences, 10(1), 24–30. https://doi.org/10.1016/j.tics.2005.11.009

Spielberg, J. M., De Leon, A. A., Bredemeier, K., Heller, W., Engels, A. S., Warren, S. L., … Miller, G. A. (2013). Anxiety type modulates immediate versus delayed engagement of attention-related brain regions. Brain and Behavior, 3(5), 532–551. https://doi.org/10.1002/brb3.157

Steinmann, S., Meier, J., Nolte, G., Engel, A. K., Leicht, G., & Mulert, C. (2017). The Callosal Relay Model of Interhemispheric Communication: New Evidence from Effective Connectivity Analysis. Brain Topography, 31(2), 218–226. https://doi.org/10.1007/s10548-017-0583-x

Techentin, C., Voyer, D., & Klein, R. M. (2009). Between- and within-ear congruency and laterality effects in an auditory semantic/emotional prosody conflict task. Brain and Cognition, 70(2), 201–208. https://doi.org/10.1016/j.bandc.2009.02.003

van Heuven, W. J. B., Mandera, P., Keuleers, E., & Brysbaert, M. (2014). Subtlex-UK: A New and Improved Word Frequency Database for British English. Quarterly Journal of Experimental Psychology, 67(6), 1176–1190. https://doi.org/10.1080/17470218.2013.850521

Vuilleumier, P. (2005). How brains beware: neural mechanisms of emotional attention. Trends in Cognitive Sciences, 9(12), 585–594. https://doi.org/10.1016/j.tics.2005.10.011

Wabnitz, P., Martens, U., & Neuner, F. (2015). Written threat: Electrophysiological evidence for an attention bias to affective words in social anxiety disorder. Cognition and Emotion, 30(3), 516–538. https://doi.org/10.1080/02699931.2015.1019837

Wambacq, I. J., & Jerger, J. F. (2004). Processing of affective prosody and lexical-semantics in spoken utterances as differentiated by event-related potentials. Cognitive Brain Research, 20(3), 427–437. https://doi.org/10.1016/j.cogbrainres.2004.03.015

Warriner, A. B., Kuperman, V., & Brysbaert, M. (2013). Norms of valence, arousal, and dominance for 13,915 English lemmas. Behavior Research Methods, 45(4), 1191–1207. https://doi.org/10.3758/s13428-012-0314-x

Watson, D., Weber, K., Assenheimer, J. S., Clark, L. A., &, et al. (1995). Testing a tripartite model: I. Evaluating the convergent and discriminant validity of anxiety and depression symptom scales. Journal of Abnormal Psychology, 104(1), 3–14. https://doi.org/10.1037//0021-843x.104.1.3

Westerhausen, R., & Hugdahl, K. (2008). The corpus callosum in dichotic listening studies of hemispheric asymmetry: A review of clinical and experimental evidence. Neuroscience & Biobehavioral Reviews, 32(5), 1044–1054. https://doi.org/10.1016/j.neubiorev.2008.04.005

Xu, Y. (2013). ProsodyPro — A Tool for Large-scale Systematic Prosody Analysis - UCL Discovery. Ucl.Ac.Uk. https://doi.org/http://discovery.ucl.ac.uk/1406070/1/Xu_TRASP2013.pdf

Xu, Y. (2019). Emotional expressions as communicative signals | Yi Xu, Andrew Kelly and Cameron Smillie. Benjamins.Com. https://doi.org/https://benjamins.com/catalog/ill.13.02xu

Zatorre, R. J. (2001). Neural Specializations for Tonal Processing. Annals of the New York Academy of Sciences, 930(1), 193–210. https://doi.org/10.1111/j.1749-6632.2001.tb05734.x

Zatorre, R. J., Belin, P., & Penhune, V. B. (2002). Structure and function of auditory cortex: music and speech. Trends in Cognitive Sciences, 6(1), 37–46. https://doi.org/10.1016/s1364-6613(00)01816-7

