## Supplement 1 for "Semantic and Prosodic Threat Processing in Trait Anxiety"

**Annex**

***Properties of Materials***

Here we report comparisons between means, to verify that both manipulations of semantic threat and prosodic threat resulted in sets of items belonging to recognizable categories. We tested differences between three acoustic properties of sentences: Hammarberg index (HI), Median Pitch (MP), Harmonicity and Shimmer. Only MP and HI were consistently different in threat vs. neutral comparisons, and consistently similar in threat vs. threat and neutral vs. neutral comparisons; where MP is higher for threatening conditions and HI lower for threatening conditions. In addition, arousal and valence were consistently different in threat vs. neutral comparisons, and consistently similar in threat vs. threat and neutral vs. neutral comparisons; where arousal is higher for threatening conditions and valence lower for threatening conditions. Figure 1 and Table 1 summarise these properties as estimates of mean differences from a Bayesian estimation supersedes the t-test (BEST) model. More details in our OSF repository (https://osf.io/z8pgf/?view_only=b5da5ce6c8644bc182231cd9b96be173).

Additionally, to confirm that sentences are perceived as threatening or neutral, we asked one group of people (n = 22) who did not take part in the main experiments to provide ratings on how threatening the 'content' of written sentences is (tapping into semantic threat), and another group (n = 10) on how threatening sentences 'sound' (tapping into acoustic threat), in both cases using 0-8 Likert scales as described in the main text. For assessing these ratings an Ordered-logistic regression was implemented. Probabilities of cutpoints (rates from 0 to 8 in the threat liker scale) are portrayed in Figure 2. These indicates that both neutral prosody and neutral semantics are clearly distinguishable from threatening prosody and semantics. However, threatening prosody is not so well distinguished as threatening semantics. Threatening prosody tend to increase by a small percentage and with greater uncertainty (see Figure 2's upper panel), which may be due to the small sample size of stimuli (just a subsample of 7 from the 54 total).

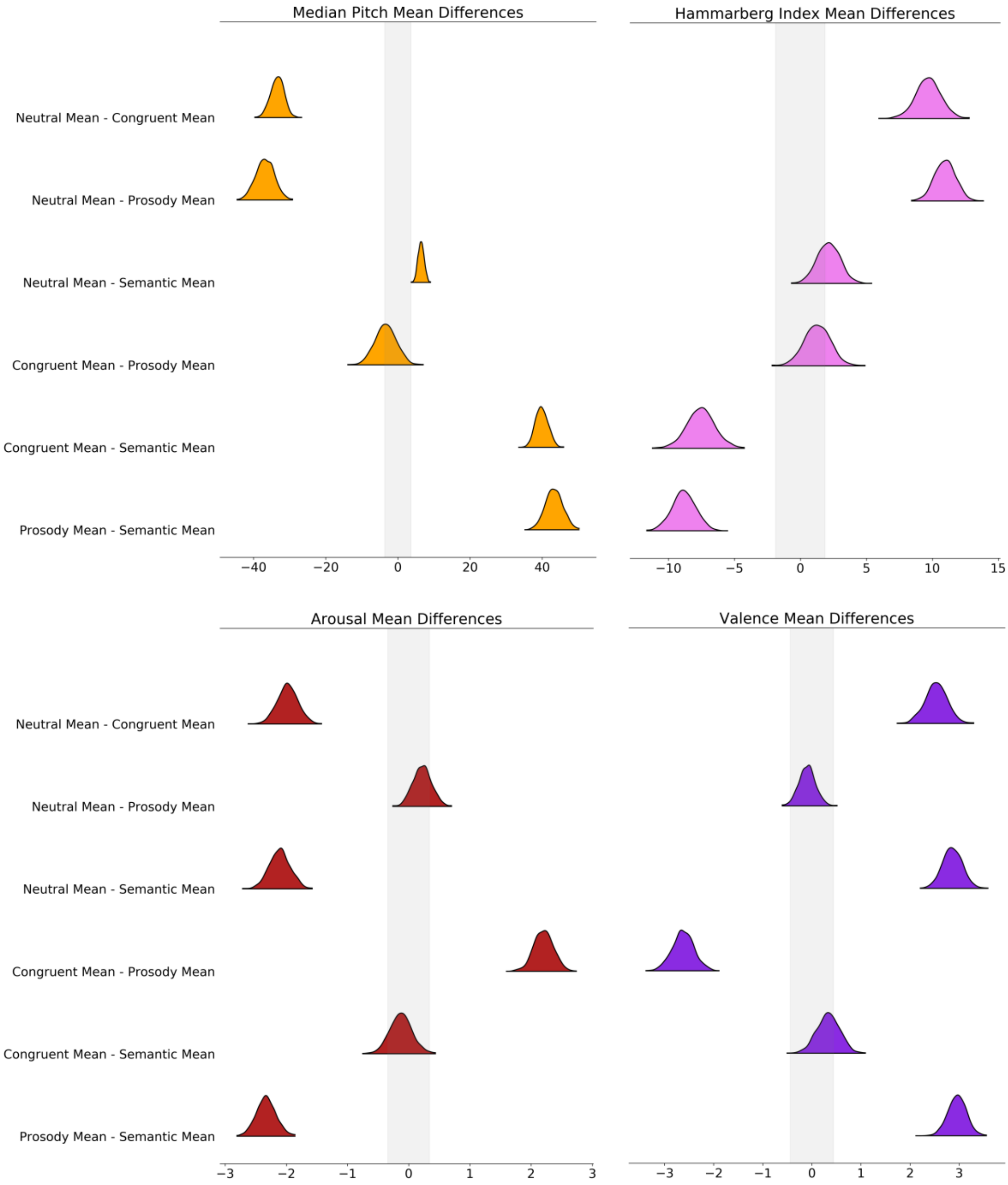


**Figure 1**. BEST estimated differences of means. Upper left: Median Pitch (MP). Lower left: arousal. Upper right: Hammarberg Index (HI). Lower right: valence. Note the similarity between MP and arousal estimates and between HI and valence estimates. This indicates that both prosodic and semantic properties of stimuli are balanced in terms of threatening and neutral speech.

Table 1. BEST differences between means and respective effect sizes (*d*)

| | Median Pitch | | | | Hammarberg Index | | | |
|---|---|---|---|---|---|---|---|---|
| **Differences** | **Mean** | **SD** | **HDI 5%** | **HDI 95%** | **Mean** | **SD** | **HDI 5%** | **HDI 95%** |
| **Neutral - Congruent** | -33.42 | 1.82 | -36.46 | -30.53 | 9.68 | 0.94 | 8.12 | 11.22 |
| **Neutral - Congruent *d*** | -4.79 | 0.61 | -5.77 | -3.84 | 2.10 | 0.25 | 1.69 | 2.52 |
| **Neutral - Prosody** | -36.81 | 2.52 | -41.32 | -33.00 | 10.96 | 0.79 | 9.73 | 12.25 |
| **Neutral - Prosody *d*** | -4.29 | 0.64 | -5.26 | -3.23 | 2.90 | 0.29 | 2.40 | 3.37 |
| **Neutral - Semantic** | 6.42 | 0.82 | 5.10 | 7.78 | 2.13 | 0.86 | 0.72 | 3.50 |
| **Neutral - Semantic *d*** | 1.98 | 0.31 | 1.47 | 2.49 | 0.52 | 0.21 | 0.19 | 0.86 |
| **Congruent - Prosody** | -3.40 | 2.94 | -7.90 | 1.69 | 1.27 | 0.99 | -0.39 | 2.80 |
| **Congruent - Prosody *d*** | -0.33 | 0.30 | -0.83 | 0.14 | 0.27 | 0.21 | -0.08 | 0.58 |
| **Congruent - Semantic** | 39.84 | 1.73 | 37.19 | 42.79 | -7.55 | 1.05 | -9.25 | -5.80 |
| **Congruent - Semantic *d*** | 6.11 | 0.80 | 4.87 | 7.38 | -1.50 | 0.24 | -1.88 | -1.09 |
| **Prosody - Semantic** | 43.24 | 2.41 | 39.27 | 47.13 | -8.83 | 0.90 | -10.23 | -7.25 |
| **Prosody - Semantic *d*** | 5.26 | 0.76 | 4.02 | 6.41 | -2.05 | 0.26 | -2.48 | -1.62 |

| | Arousal | | | | Valence | | | |
|---|---|---|---|---|---|---|---|---|
| **Differences** | **Mean** | **SD** | **HDI 5%** | **HDI 95%** | **Mean** | **SD** | **HDI 5%** | **HDI 95%** |
| **Neutral - Congruent** | -1.98 | 0.17 | -2.28 | -1.73 | 2.53 | 0.22 | 2.12 | 2.87 |
| **Neutral - Congruent *d*** | -2.35 | 0.30 | -2.82 | -1.86 | 2.33 | 0.30 | 1.84 | 2.82 |
| **Neutral - Prosody** | 0.22 | 0.15 | 0.00 | 0.49 | -0.10 | 0.17 | -0.38 | 0.17 |
| **Neutral - Prosody *d*** | 0.32 | 0.22 | -0.03 | 0.68 | -0.11 | 0.19 | -0.45 | 0.20 |
| **Neutral - Semantic** | -2.11 | 0.17 | -2.39 | -1.83 | 2.86 | 0.20 | 2.54 | 3.18 |
| **Neutral - Semantic *d*** | -2.54 | 0.33 | -3.02 | -1.94 | 2.88 | 0.32 | 2.35 | 3.40 |
| **Congruent - Prosody** | 2.20 | 0.16 | 1.97 | 2.48 | -2.63 | 0.22 | -3.03 | -2.30 |
| **Congruent - Prosody *d*** | 2.74 | 0.32 | 2.21 | 3.26 | -2.52 | 0.32 | -3.06 | -2.05 |
| **Congruent - Semantic** | -0.13 | 0.18 | -0.41 | 0.19 | 0.33 | 0.23 | -0.01 | 0.73 |
| **Congruent - Semantic *d*** | -0.14 | 0.19 | -0.45 | 0.19 | 0.28 | 0.20 | -0.03 | 0.61 |
| **Prosody - Semantic** | -2.33 | 0.16 | -2.59 | -2.06 | 2.96 | 0.19 | 2.63 | 3.24 |
| **Prosody - Semantic *d*** | -2.94 | 0.34 | -3.45 | -2.35 | 3.14 | 0.34 | 2.57 | 3.67 |

***Note:*** *All estimates are from posterior distributions of differences between means and effect sizes.*

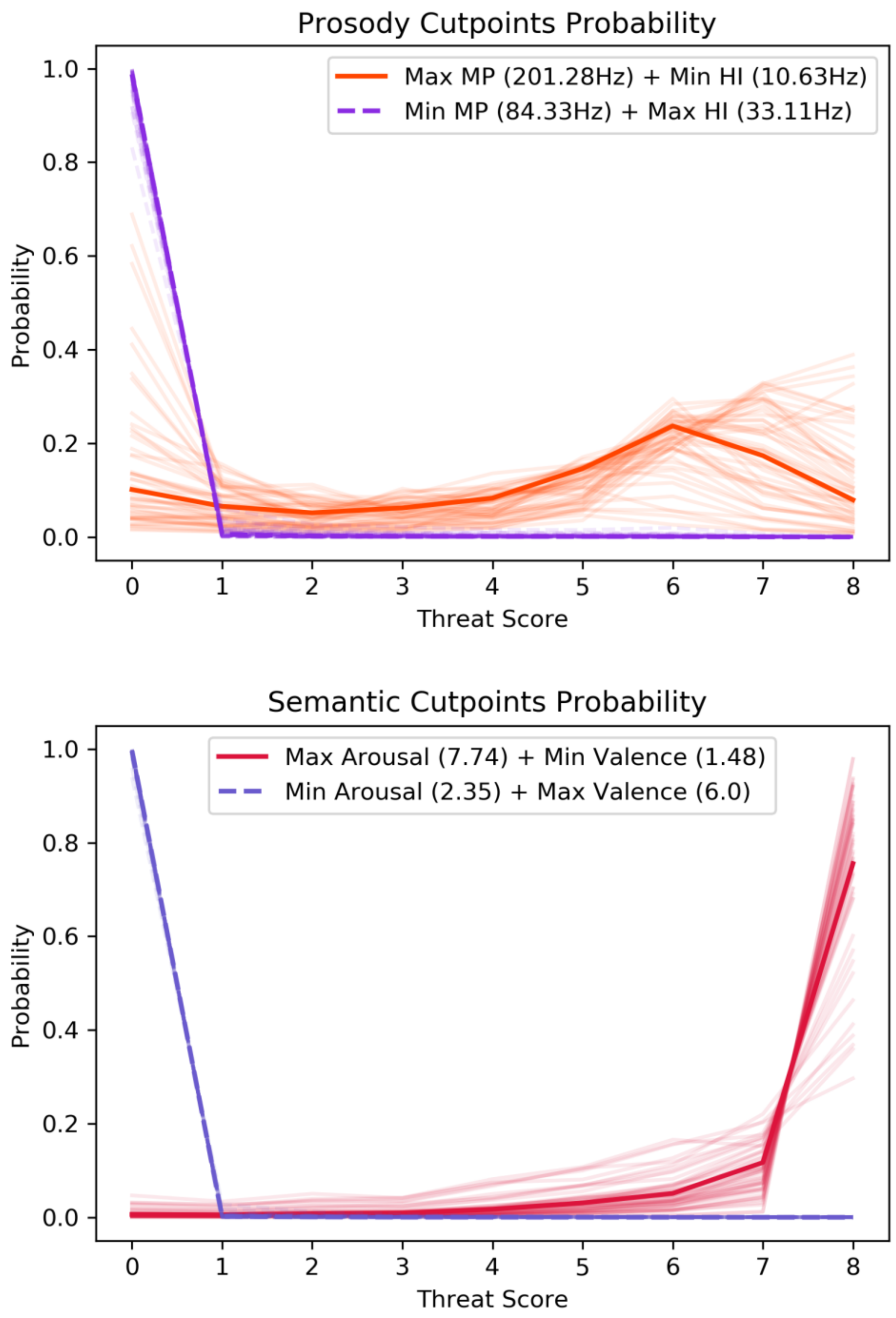


**Figure 2**. Ordered-logistic regression cutpoints estimates. Up: probability of rating threat (0 to 8) when listening to Prosody stimuli (prosodic threat). Bottom: probability of rating threat (0 to 8) when listening to Semantic stimuli (semantic threat). Lines indicate posterior means, faded lines indicate uncertainty (random samples from posterior). Note that when median pitch (MP) is the maximum and Hammarberg index (HI) is at the minimum, the probability of rating sentences as 5 or above in threat scale goes up by near 10%, a consistent but somewhat uncertain and small increase. However, when MP is at the minimum and HI at maximum, the probability of rating a sentence as 1 or below in the threat scale goes up near 100%. When arousal is at maximum and valence at minimum, the probability of rating a sentence over 7 points increases near 70%; and when arousal is at minimum and valence at maximum the probability of rating 1 or below goes up to near 100%.
